# Co-culture and biogeography of *Prochlorococcus* and SAR11

**DOI:** 10.1101/460428

**Authors:** Jamie W. Becker, Shane L. Hogle, Kali Rosendo, Sallie W. Chisholm

**Affiliations:** Department of Civil and Environmental Engineering, Massachusetts Institute of Technology, Cambridge, MA, USA; Department of Biology, Massachusetts Institute of Technology, Cambridge, MA, USA

## Abstract

*Prochlorococcus* and SAR11 are among the smallest and most abundant organisms on Earth. With a combined global population of about 2.7 x 10^28^ cells, they numerically dominate bacterioplankton communities in oligotrophic ocean gyres and yet they have never been grown together *in vitro*. Here we describe co-cultures of *Prochlorococcus* and SAR11 isolates representing both high- and low-light adapted clades. We examined: (1) the influence of *Prochlorococcus* on the growth of SAR11 and *vice-versa*, (2) whether *Prochlorococcus* can meet specific nutrient requirements of SAR11, and (3) how co-culture dynamics vary when *Prochlorococcus* is grown with SAR11 compared with sympatric copiotrophic bacteria. SAR11 grew as much as 70% faster in the presence of *Prochlorococcus*, while the growth of the latter was unaffected. When *Prochlorococcus* populations entered stationary phase, SAR11 abundances decreased dramatically. In parallel experiments with copiotrophic bacteria however, the heterotrophic partner increased in abundance as *Prochlorococcus* densities leveled off. The presence of *Prochlorococcus* was able to meet SAR11’s central requirement for organic carbon, but not reduced sulfur. *Prochlorococcus* strain MIT9313, but not MED4, could meet the unique glycine requirement of SAR11, likely due to production and release of glycine betaine by MIT9313. Evidence suggests that *Prochlorococcus* MIT9313 may also compete with SAR11 for the uptake of 3-dimethylsulfoniopropionate (DMSP). To place our results in context, we assessed the relative contribution of *Prochlorococcus* and SAR11 genome equivalents to those of identifiable bacteria and archaea in over 800 marine metagenomes. At many locations, more than half of the identifiable genome equivalents in the euphotic zone belonged to *Prochlorococcus* and SAR11 – highlighting the biogeochemical potential of these two groups.

## Introduction

The global ocean is numerically dominated by *Prochlorococcus* and SAR11 (*Pelagibacterales*). *Prochlorococcus*, a cyanobacterium, is the most abundant primary producer in tropical and subtropical waters, where its estimated 2.9 x 10^27^ cells produce ca. 4 Gt of organic carbon annually [1]. As such, they support a notable fraction of the secondary production in these nutrient-poor waters [2]. Members of the alphaproteobacteria known as SAR11 are found throughout the marine environment with an estimated global abundance of 2.4 x 10^28^ cells, about half of which are in the euphotic zone [3]. *Prochlorococcus* and SAR11 together have been estimated to comprise roughly 40-60% of the total bacteria in oligotrophic surface waters at Station ALOHA in the North Pacific [4].

Since its discovery three decades ago, *Prochlorococcus* has emerged as a powerful model organism for microbial ecology [5]. Its ecotype diversity is well-characterized [6-9] and there is an increasing wealth of genomic, transcriptomic and proteomic information applied to this group [10-12]. The discovery of SAR11 a few years later further advanced our understanding of microbial ecology and evolution in the oligotrophic ocean [13,14]. SAR11 possesses many traits that highlight adaptations to the oligotrophic marine environment, including a small genome size [15,16] and unique metabolic dependencies [17-20] partitioned among diverse ecotypes with distinct biogeography [21-24].

In addition to having large population sizes consisting of genetically distinct ecotypes, or clades, *Prochlorococcus* and SAR11 have adapted to their oligotrophic habitat by minimizing their genome content and metabolic versatility. SAR11 and high-light adapted *Prochlorococcus* have small cell sizes between 0.2 and 0.8 microns in diameter, small genomes (1.2 to 1.8 Mb) with low guanine-cytosine content (29 to 32 %GC) [10,15], and fewer regulatory σ-factors than would be predicted from the size of their genomes [25]. Low-light adapted *Prochlorococcus* genomes are larger (ca. 2.5 Mb) with higher guanine-cytosine content (33-50 %GC), but are still relatively streamlined compared to copiotrophic bacteria. Genome reduction has resulted in the loss of metabolic capabilities in both *Prochlorococcus* and SAR11, some of which, such as reduction of oxidative stress [26] or a requirement for exogenous reduced sulfur [17] respectively, are provided by neighboring microbes. Likely as a result of these types of interdependencies, both taxa are difficult to isolate and maintain *in-vitro* compared to faster-growing (r-selected) microbes; many of the same traits that confer a competitive advantage in their native habitat likely render them difficult to maintain in a laboratory.

*Prochlorococcus* and SAR11 cells can interact with each other in the surface ocean either directly or indirectly through the exchange of metabolites. Like all heterotrophs, SAR11 cells rely on organic matter derived from primary producers and recent experiments revealed that some members of the SAR11 clade likely rely heavily on organic compounds as an important source of phosphorus in addition to carbon [27]. SAR11 also has several unique metabolic requirements that implicate potential mutualism with *Prochlorococcus*. For example, SAR11 exhibits auxotrophy for a thiamin precursor molecule found in spent medium from *Prochlorococcus* [20] and some strains have a glycine requirement that can be partially met by glycolate, a byproduct of *Prochlorococcus* photorespiration [2,28]. Furthermore, evidence from comparative genomics and metabolic modeling suggests that *Prochlorococcus* and SAR11 may exchange glycolate, pyruvate, and malate through complementary and/or co-dependent metabolic pathways, and it is likely that the evolution of these two organisms has been tightly interwoven [29].

Despite the high abundances and frequently observed habitat overlap of *Prochlorococcus* and SAR11 in the tropical and subtropical oligotrophic ocean, a system for co-culturing and exploring interactions between members of these two model groups has remained elusive, largely due to their cryptic growth requirements and inherent challenges in maintaining laboratory isolates. Here we report the development of stable (over two year) co-cultures of *Prochlorococcus* and SAR11, and explorations into the interrelated growth dynamics of these groups. Specifically, we sought to determine if *Prochlorococcus* can provide SAR11 with individual growth requirements and how SAR11/*Prochlorococcus* co-culture dynamics compare with the co-culture of *Prochlorococcus* and a suite of sympatric copiotrophic bacteria. Given the high abundance of these two groups throughout the oligotrophic surface ocean, these co-cultures establish a novel and ecologically significant model system for the study of marine microbial ecology.

## Materials and methods

### Biogeography analysis

The computational steps for determining the genome equivalents of SAR11, *Prochlorococcus* and other identifiable bacteria and archaea are described in detail in the supplementary methods. Briefly, we analyzed 195 surface, mixed layer, and deep chlorophyll maximum layer metagenomes from the Tara oceans project [30-32], 480 metagenome samples acquired during GEOTRACES cruises, and 133 metagenome samples from the HOT and BATS oceanographic time-series [33]. Metagenome reads were Illumina adapter trimmed, quality filtered, and overlapped using the bbtools (BBMap V37.90) software suite [34]. We annotated the quality controlled metagenomes against a custom reference database of approximately 26 000 bacterial, archaeal, viral, and microbial eukaryotic isolate, single cell, and metagenome and transcriptome assembled genomes compiled from various sources [35-43] using Kaiju (V1.6.0; [44]). The taxonomic composition of the reference database was intended to predominantly reflect that of the marine environment, while minimizing (but not excluding) the representation of clinical, industrial, and terrestrial host-associated genomes/samples. The majority of reads in our study (54% ± 10%) could be classified across all metagenomes using this approach (see supplementary methods).

We extracted the reads from each metagenome classified by Kaiju as *Prochlorococcus* (genus), SAR11 (order *Pelagibacterales*), and bacteria/archaea (kingdom), and then quantified universal, single-copy marker genes within each taxonomically resolved pool using MicrobeCensus (V1.1.1; [45]). The abundances of these marker genes were used to estimate ‘genome equivalents’ (the operational number of genomes represented by single copy marker genes) within each taxonomically resolved read pool. We report abundances as marker-gene resolved genome equivalents rather than total classified reads due to systematic variations in the average genome size between groups like *Prochlorococcus* and SAR11 and the rest of the microbial community [45]. Here, the relative abundance of *Prochlorococcus* or SAR11 is defined by the fraction of *Prochlorococcus* or SAR11 genome equivalents divided by the total number of bacterial and archaeal genome equivalents that could be identified in each sample. Phylogenies presented in the supplemental material are derived from a concatenated protein multiple sequence alignment based on 120 taxonomically conserved single copy marker genes [41,46] and were inferred with RAxML (V8.2.9; [47]).

### Strain selection and isolation

*Prochlorococcus* strains were chosen to represent clades with genetic and physiological distinctions that influence their biogeographic distributions [10,48] with a preference for strains isolated from the N. Atlantic Ocean, the place of origin of *Pelagibacterales sp.* HTCC7211 (Table 1). HTCC7211 was isolated in 2006 from 10 m at the Bermuda Atlantic Time-series Study site in the Sargasso Sea and is a member of the abundant warm-water surface-dwelling Ia.3 ecotype [24] (Fig. S1). Several new strains of *Prochlorococcus* and heterotrophic bacteria were isolated to examine interactions between isolates from the same water sample – i.e. sympatric strains where the ‘chain of custody’ was documented. Details regarding their isolation are included in Table 1 and supplementary methods.

**Table 1.** Isolates used in this study. The phylogenetic affiliations of the *Prochlorococcus* and SAR11 strains are shown in Fig. S1.

| *Prochlorococcus* |  |  |  |  |  |
| --- | --- | --- | --- | --- | --- |
| Strain | Clade | Isolation Location | Depth (m) | Year | Reference |
| MED4 | HLI | Mediterranean Sea | 5 | 1989 | [1] |
| MIT9312 | HLII | Gulf Stream | 135 | 1993 | [2] |
| MIT9301 | HLII | Sargasso Sea (BATS) | 90 | 1993 | [3] |
| MIT1314 | HLII | North Pacific (Station ALOHA) | 150 | 2013 | This study |
| MIT0801 | LLI | Sargasso Sea (BATS) | 40 | 2008 | [4] |
| MIT9313 | LLIV | Gulf Stream | 135 | 1993 | [2] |
| MIT1320 | LLIV | North Pacific (Station ALOHA) | 150 | 2013 | [5] |
| MIT1327 | LLIV | North Pacific (Station ALOHA) | 150 | 2013 | [5] |
| **Heterotrophic bacteria** |  |  |  |  |  |
| Strain | Order (clade/genus) | Isolation Location | Depth (m) | Year | Reference |
| HTCC7211 | SAR11 (*Pelagibacterales*) | Sargasso Sea (BATS) | 10 | 2006 | [6] |
| MIT1351 | *Rhodospirillales* (*Thalassospira*) | North Pacific (Station ALOHA) | 150 | 2013 | This study |
| MIT1352 | *Rhodobacterales* (*Roseobacter*) | North Pacific (Station ALOHA) | 150 | 2013 | This study |
| MIT1353 | *Alteromonadales* (*Marinobacter*) | North Pacific (Station ALOHA) | 150 | 2013 | This study |

1. Chisholm SW, Frankel SL, Goericke R, Olson RJ, Palenik B, Waterbury JB, et al. *Prochlorococcus marinus* nov. gen. nov. sp.: an oxyphototrophic marine prokaryote containing divinyl chlorophyll a and b. *Arch. Microbiol.* 1992;157:297–300. 2. Moore LR, Rocap G, Chisholm SW. Physiology and molecular phylogeny of coexisting *Prochlorococcus* ecotypes. *Nature* 1998;393:464–7. 3. Rocap G. Genetic diversity and ecotypic differentiation in the marine cyanobacteria *Prochlorococcus* and *Synechococcus*. PhD [dissertation]. Cambridge (MA): Massachusetts Institute of Technology; 2000. 4. Biller SJ, Berube PM, Berta-Thompson JW, Kelly L, Roggensack SE, Awad L, et al. Genomes of diverse isolates of the marine cyanobacterium *Prochlorococcus*. *Sci. Data* 2014;1:140034–11. 5. Cubillos-Ruiz A, Berta-Thompson JW, Becker JW, van der Donk WA, Chisholm SW. Evolutionary radiation of lanthipeptides in marine cyanobacteria. *Proc. Natl. Acad. Sci. U.S.A.* 2017;114:E5424–33. 6. Stingl U, Tripp HJ, Giovannoni SJ. Improvements of high-throughput culturing yielded novel SAR11 strains and other abundant marine bacteria from the Oregon coast and the Bermuda Atlantic Time Series study site. *ISME J* 2007;1:361–71.

### Development of ProMS medium for growing both *Prochlorococcus* and SAR11

While media recipes exist for culturing both *Prochlorococcus* [49] and SAR11 [28,50], none that we tested could support the growth of both strains. To this end, we established ProMS, a medium with a 0.2 μm filtered, autoclaved Sargasso surface seawater base, that was capable of supporting the growth of each strain in both mono -and co-culture (Table S1). After autoclaving, the seawater was sparged with sterile CO_2_ followed by air to reestablish a bicarbonate-based buffer system [51] and amended with sterile Pro99 nutrients [49] and organic compounds to meet the known unique metabolic requirements of SAR11 [52] as follows: pyruvate (1 μM), glycine (1 μM), methionine (0.2 μM), and the vitamin mix developed for AMS1 medium [28] diluted 50-fold. Organic additions were modeled after the medium developed by Carini et al. (2013), with reduced concentrations designed to promote interactions.

In preparation for co-culture experiments, axenic SAR11 cells were first transferred from AMS1 medium amended with pyruvate (50 μM), glycine (50 μM) and methionine (10 μM) to ProMS medium. Prior to co-culturing experiments, monocultures of *Prochlorococcus* and SAR11 were maintained using ProMS medium in acid-washed autoclaved polycarbonate tubes at 22 ° C under constant illumination (12 μmol photons m^−2^ s^−1^) for >25 consecutive transfers (>100 generations).

### Co-cultures of *Prochlorococcus* and SAR11

Following acclimation of individual strains to ProMS medium, replicate batch co-cultures of SAR11 and diverse strains of *Prochlorococcus* were established at an initial ratio of 10:1 *Prochlorococcus*:SAR11 in acid-washed autoclaved polycarbonate tubes. Co-cultures were monitored for growth and purity by flow cytometry until both populations entered stationary and/or death phase.

### Organic nutrient substitution experiments

Monocultures of SAR11 and co-cultures of SAR11 and *Prochlorococcus* strains MED4 and MIT9313 were conditioned (where possible) onto versions of ProMS medium lacking either pyruvate, glycine or methionine to examine the ability of *Prochlorococcus* to supply SAR11 with specific nutrient requirements. SAR11 exhibited no change in maximum cell abundance in ProMS lacking pyruvate (likely due to the availability of compounds present in the Sargasso seawater base that can meet the central carbon requirement of SAR11), so the concentrations of glycine, methionine and vitamins in ProMS were increased 50-fold to create ProMC medium and induce pyruvate limitation.

### Co-cultures of *Prochlorococcus* and sympatric copiotrophs

For experiments with sympatric *Prochlorococcus* and heterotrophs, following acclimation of axenic strains to Pro99 medium, triplicate pairwise batch co-cultures of *Prochlorococcus* strains MIT1314 and MIT1327 and heterotrophic bacteria strains MIT1351, MIT1352 and MIT1353 were established at an initial ratio of either 150:1 or 300:1 *Prochlorococcus*:heterotrophic bacteria for co-cultures involving MIT1314 and MIT1327 respectively (Table 1). Mono -and co-cultures were maintained in acid-washed autoclaved borosilicate glass tubes and monitored for growth and purity as described above.

### Enumeration of cells

Cell concentrations were determined using a Guava Technologies easyCyte 12HT flow cytometer (EMD Millipore) after staining with SYBR green I (Lonza) in the dark for at least 55 min. Samples were diluted in sterilized Sargasso seawater to ensure <500 cells μl^−1^ to avoid coincidence counting. Samples were run with only the blue (488 nm) excitation laser enabled for maximum power and populations were resolved based on their green (525/30) and red (695/50) emission parameters.

## Results and Discussion

### Co-occurrence of *Prochlorococcus* and SAR11 in the wild

We leveraged the recent expansion of marine metagenomic data to perform a global census of the *Prochlorococcus* genus and the *Pelagibacterales* order (hereafter SAR11) by estimating their relative abundance throughout the global ocean. We examined 668 shotgun metagenomes with a broad geographic distribution collected within the euphotic zone [30-33] and 133 metagenomes from the Bermuda Atlantic Time-series (BATS) and Hawaii Ocean Time-series (HOT) [33]. Total metagenome recruitment combined with the enumeration of single copy marker genes was used to estimate the contribution of each group to the total identifiable bacterial and archaeal genome equivalents in each sample (see supplementary methods for additional information).

The relative abundance of SAR11 ranged between 7% and 55% (Fig. 1). Their contribution was largely horizontally and vertically consistent in the euphotic zone, although somewhat lower at high latitudes and coastal sampling sites, and in deeper waters near the base of this zone (Figs. 1 and S3). In contrast, the distribution of *Prochlorococcus* was bounded by 45 degrees N/S; its relative abundance began to decline sharply near 40 degrees N/S, consistent with previous reports using other approaches [1,8]. The relative abundance of *Prochlorococcus* ranged from undetectable to >45% in parts of the remote south Pacific Ocean (Fig. 1). The contribution of *Prochlorococcus* generally decreased with depth and light intensity and its maximum was typically found in the upper 100 meters (Fig. S3).

**Figure 1.**
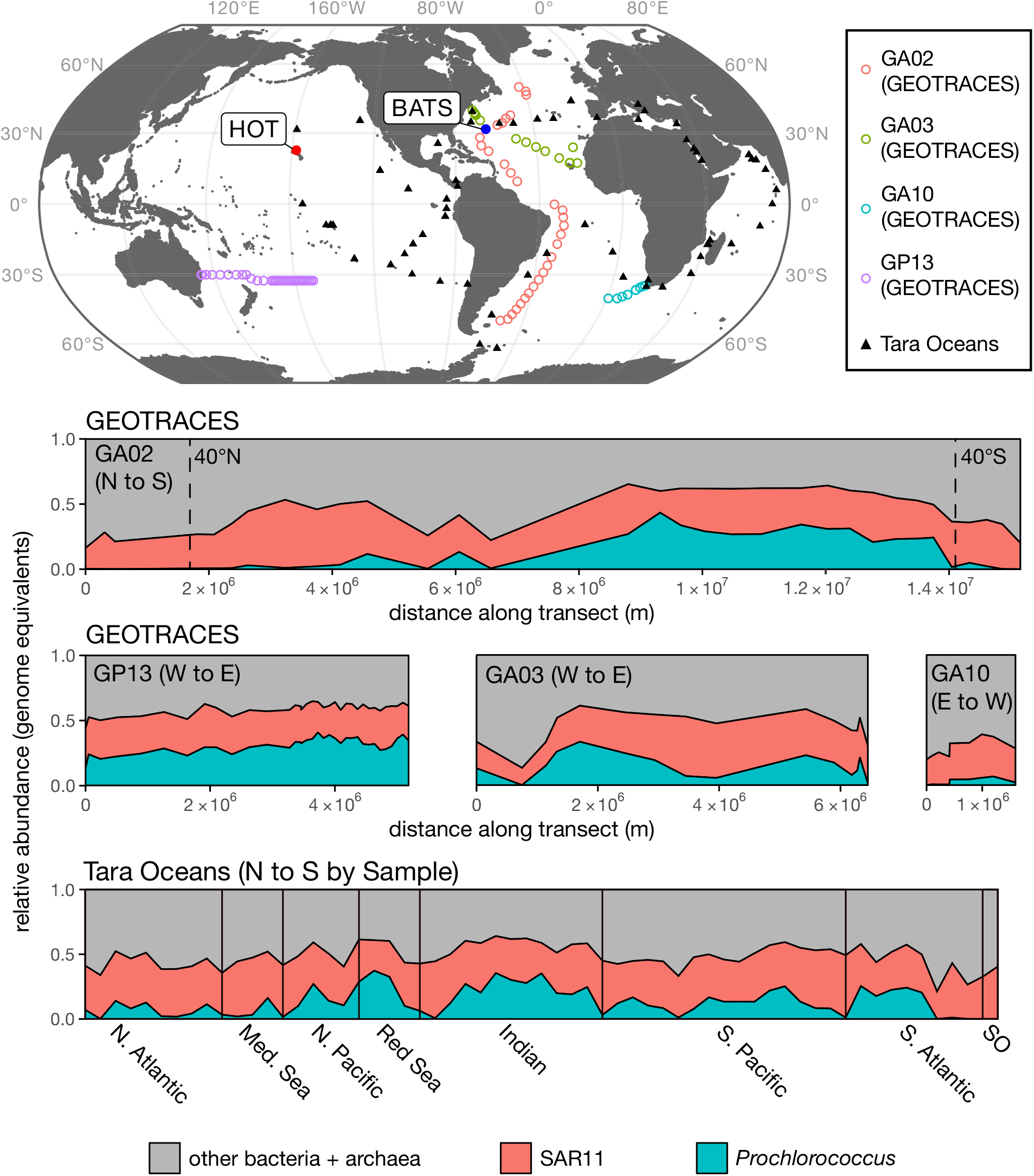
Genome equivalents of *Prochlorococcus* and SAR11 relative to total identifiable bacteria and archaea in the surface ocean. (Upper) Locations of GEOTRACES, HOT and BATS Time-series, and Tara Oceans metagenome samples used for the analysis. (Lower) Vertical axes represent the abundance of *Prochlorococcus* (teal) and SAR11 (red) genome equivalents relative to all other identifiable bacteria and archaea (gray) throughout the upper 50 meters of the global ocean. GEOTRACES horizontal axes depict the distance along each transect, while the Tara Oceans horizontal axis displays evenly spaced samples organized by latitude (N to S) within each oceanic region. Forty degrees N/S are marked on the GA02 panel.

Both *Prochlorococcus* and SAR11 are known to display seasonal growth dynamics [7,53] – also evident here in metagenomes from the HOT and BATS time-series sites (Fig. 2). In surface waters and the deep chlorophyll maximum (DCM) layer at the BATS site, the relative abundance of *Prochlorococcus* varied from nearly undetectable during the spring, to over 20% of the identifiable bacterial and archaeal community in late autumn, as the depth of the mixed layer increased – consistent with previous reports [7]. In contrast, the relative abundance of SAR11 peaked about when the mixed layer depth shoaled to its most shallow depth, consistent with previously reported patterns for members of the SAR11 Ia clade [23]. The relative abundances of *Prochlorococcus* and SAR11 displayed less seasonality in surface waters at station ALOHA, but did undergo sporadic oscillations (e.g. June 2003) that may reflect mesoscale events such as nutrient influxes, temperature shifts, water transport, or entrainment caused by eddies [54]. In summary, the combined genome equivalents of *Prochlorococcus* and SAR11 regularly comprised more than half of the total identifiable bacteria and archaea in the euphotic zone of the oligotrophic tropical and subtropical ocean. We stress however, that these genome equivalents do not equate with cell concentrations, as some cells may have multiple copies of their genomes, and furthermore, should not be misinterpreted as representing relative biomass, as these are among the smallest bacteria in the marine ecosystem.

**Figure 2.**
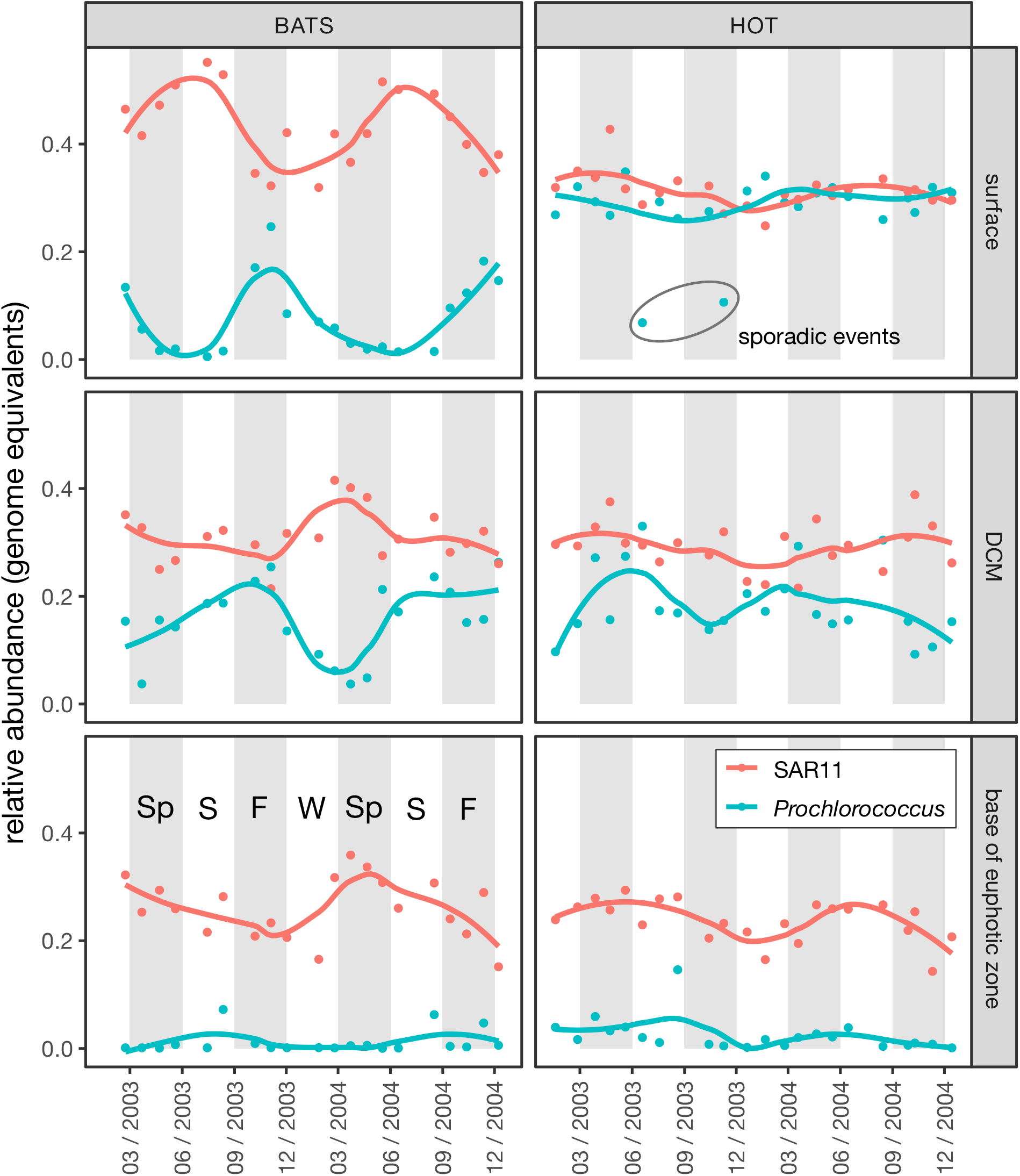
Seasonality of *Prochlorococcus* (teal) and SAR11 (red) estimated genome equivalents in metagenomes sampled from two ocean time-series sites - the Bermuda Atlantic Time-series (BATS) and Hawaii Ocean Time-series (HOT). Vertical axes represent the abundance of each group (genome equivalents) relative to total identifiable bacteria and archaea genome equivalents, while horizontal axes represent time (month/year). Data are faceted by time series (horizontal) and depth (vertical). Surface samples are from within the mixed layer (≤ 25m), DCM corresponds to the depth of maximum chlorophyll a fluorescence, and the base of the euphotic zone is defined by the depth at which ca. 1% of surface photosynthetically active radiation remains (see [33] for details). Solid lines represent a local weighted regression analysis (LOESS) smooth function for each taxonomic group at each depth. Anomalously low abundances of *Prochlorococcus* are denoted as sporadic events.

### Co-culture of *Prochlorococcus* and SAR11

Motivated by the overlapping biogeography of *Prochlorococcus* and SAR11 and their near ubiquity and high relative abundances in the surface ocean, we explored interactions between members of these groups by developing co-cultures of *Pelagibacterales sp.* HTCC7211 and a number of *Prochlorococcus* strains representing high-light and low-light adapted clades (Table 1). HTCC7211 (hereafter “SAR11”) was isolated from surface waters in the Sargasso Sea [55], making it an ideal candidate for co-culture with several of the *Prochlorococcus* strains isolated from the same location (Table 1). Through many rounds of trial and error, we designed a natural seawater-based medium (ProMS) that could support the growth of both cell types independently (Table S1). The growth rate of SAR11 increased from 0.35 ± 0.01 d^−1^ when grown in AMS1 medium to 0.41 ± 0.01 d^−1^ after acclimation to ProMS, however the maximum cell abundance decreased from 1.1 x 10^8^ cells ml^−1^ to 7.5 x 10^6^ cells ml^−1^ (Fig. S2) due to the decrease in organic nutrient concentrations. The transfer of axenic *Prochlorococcus* from Pro99 to ProMS medium did not change its growth rate or maximum cell abundance.

We then propagated the strains in semi-continuous co-culture, where they remained stable for at least 2 years (Fig. 3). The frequency and magnitude of culture dilution was determined from experience, based on the cell concentrations of *Prochlorococcus*: if they exceeded ca. 5 x 10^8^ cells ml^−1^, the culture would transition into stationary phase and the SAR11 population would decline rapidly (see below). If diluted below 10^6^ cells ml^−1^, *Prochlorococcus* would display a lag phase or not grow at all – especially low-light adapted strains. Once stabilized within these limits, *Prochlorococcus* remained in log-phase growth if diluted to 1.25 x 10^6^ cells ml^−1^ every 3 to 4 days. Because dilution metrics were dictated solely by the density of *Prochlorococcus*, there was no reason *a priori* that SAR11 abundances would remain in quasi-steady state. At monoculture growth rates, SAR11 densities would have been diluted down to <1 cell ml^−1^ within three months using these dilution rates. That they reached a quasi-steady state when grown with *Prochlorococcus* implies that the metabolisms of the autotroph and heterotroph became coupled in some way, resulting in a relatively constant SAR11 to *Prochlorococcus* cell ratio of 1:3 throughout the co-culture period.

**Figure 3.**
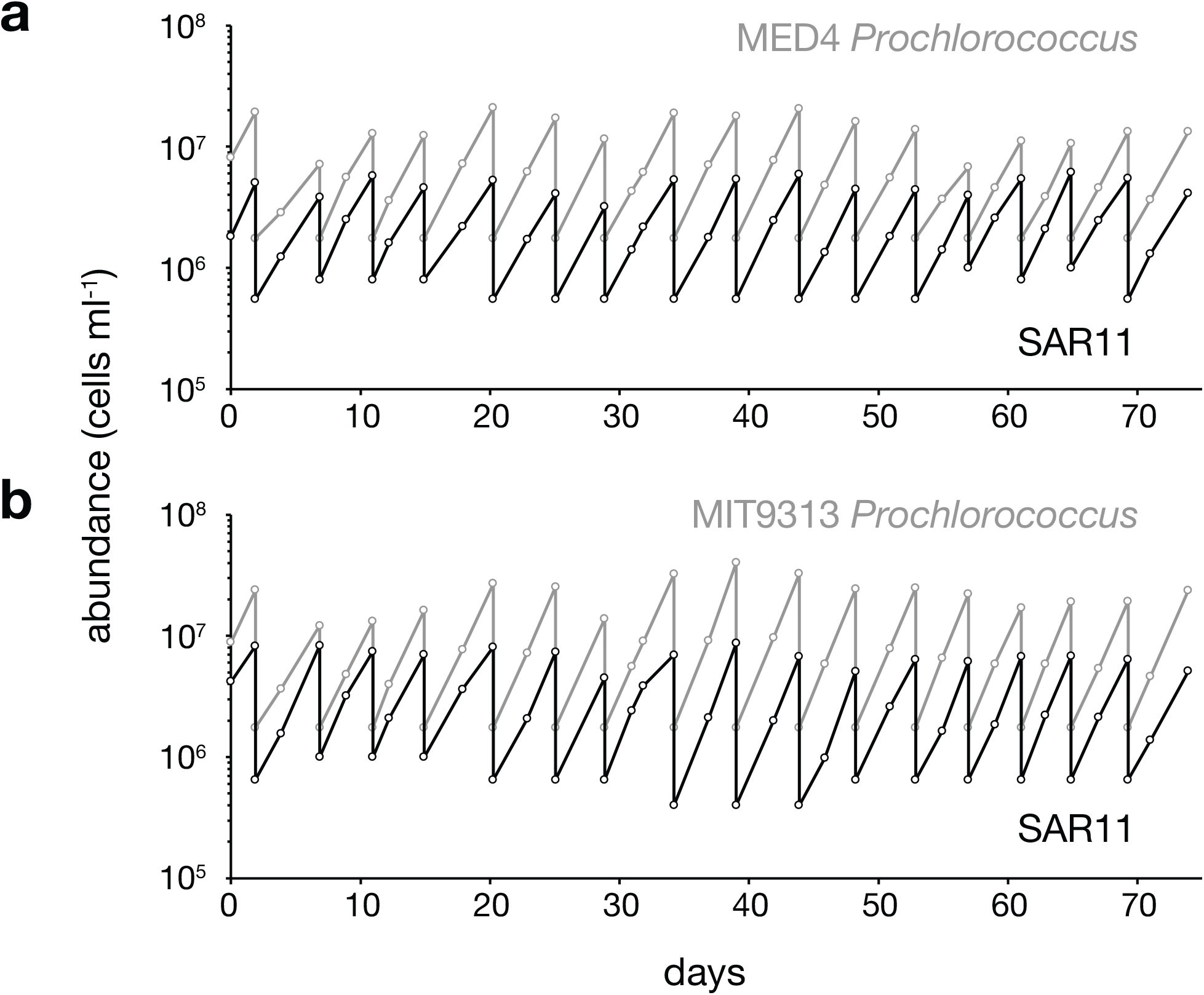
Cell abundance as a function of time in log-phase, semi-continuous batch co-cultures of SAR11 (*Pelagibacterales sp.* HTCC7211) (black lines) and *Prochlorococcus* strain MED4 (a; gray line) and MIT9313 (b; gray line) in ProMS medium. Dilution frequency and volume were dictated by *Prochlorococcus* cell density (see text).

### *Prochlorococcus* and SAR11 growth dynamics in batch co-cultures

To examine these interactions in detail, SAR11 was co-cultured with 4 strains of *Prochlorococcus* representing two high-light adapted (HL) and two low-light adapted (LL) clades (Fig. S1, Table 1). SAR11 and *Prochlorococcus* cells were taken from early log-phase axenic cultures, inoculated into co-culture, and their growth patterns were monitored in Mono -and co-culture. The growth of *Prochlorococcus* was not significantly influenced by the presence of SAR11 over the entire growth curve (Fig. S4). SAR11 however, grew 15 to 70% faster (depending on the *Prochlorococcus* strain) in the co-cultures than it did when grown alone (Fig. 4). SAR11 always entered stationary phase earlier than *Prochlorococcus* in the co-cultures, reaching maximum cell abundances that were at, or slightly above (up to 2-fold higher) those attained in monoculture (Fig. 4).

**Figure 4.**
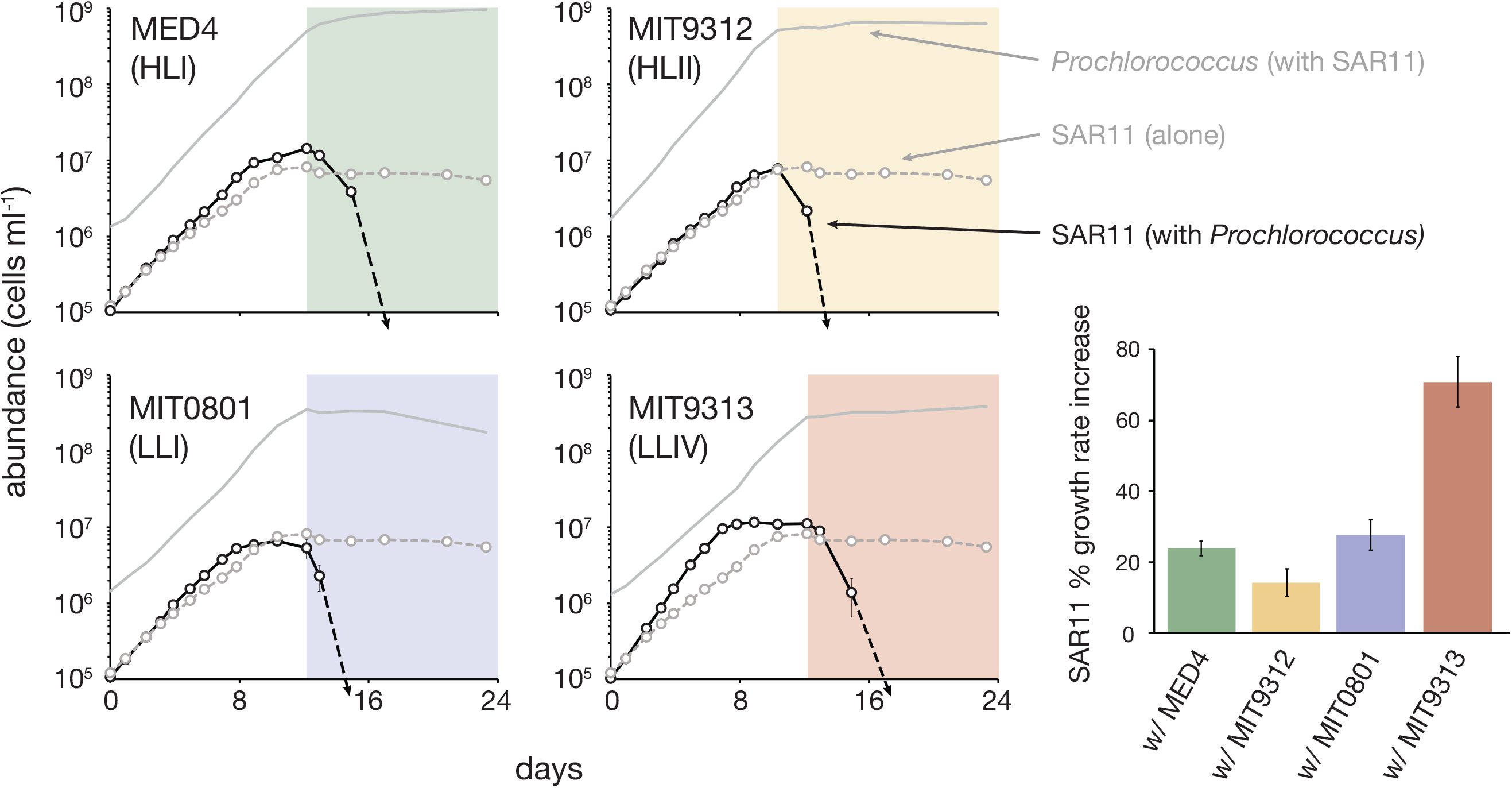
Growth of *Prochlorococcus* (solid gray lines, reproduced from Fig. S4) and SAR11 (*Pelagibacterales sp.* HTCC7211) (black lines) in co-culture compared to the growth of SAR11 monocultures (dashed gray lines). Growth curves for *Prochlorococcus* monocultures are presented in Fig. S4. Shaded regions denote the interval when *Prochlorococcus* are in stationary phase. SAR11 cells were undetectable in co-culture with *Prochlorococcus* after day 12 to 15 (dashed black arrows). Bar graph on the right depicts the percent increase in SAR11 growth rate due to the presence of each *Prochlorococcus* strain. The growth rate of SAR11 alone was 0.41 day-1. Circles and bars represent the mean (± s.d.) of biological triplicates. Error bars are smaller than the size of the symbols where not visible.

Precisely when *Prochlorococcus* entered stationary phase (beginning of colored shading in Fig. 4), SAR11 cell concentrations declined abruptly to below detection limits – in striking contrast to its behavior when grown alone where cell numbers simply level off in stationary phase (Fig. 4). The antagonistic effect of the co-culture conditions on SAR11 was apparent for all *Prochlorococcus* strains, revealing a condition-dependent shift likely caused by a growth phase-specific release of metabolites from *Prochlorococcus*, as has been observed in eukaryotic phytoplankton [56,57]. Certain amino acids and osmolytes at high concentrations have been shown to slow or even prohibit the growth of this SAR11 strain [18,58], which may explain our observations.

### *Prochlorococcus* co-cultured with sympatric copiotrophic bacteria

Although all but one of the *Prochlorococcus* strains discussed above were isolated from the same ocean as the SAR11 strain, they were not isolated from the same water sample; our attempts to isolate truly sympatric *Prochlorococcus* and SAR11 failed. We did, however, isolate two new *Prochlorococcus* strains (MIT1314 - HLII clade, and MIT1327 - LLIV clade, see Fig. S1, Table 1) along with a number of sympatric copiotrophic heterotrophic bacteria – i.e. strains that grow rapidly in media rich in organic matter. These heterotrophic strains have genomic characteristics (4.4 Mb; 55 - 59 %GC) typical of copiotrophic bacteria [59] and represent taxa found at low abundances in the oligotrophic ocean, except when subjected to episodic disturbances [60]. We studied 3 of these – *Thalassospira sp.* MIT1351, *Roseobacter sp.* MIT1352, and *Marinobacter sp*. MIT1353 –in co-culture with sympatric *Prochlorococcus* to compare with SAR11 patterns (Table 1).

In contrast to SAR11, which would not grow alone in Pro99 medium [49] unless amended with labile substrates to meet its unique metabolic requirements (see Methods, Table S1), the copiotrophic strains grew appreciably in unamended Pro99 medium, apparently using their more diverse metabolic repertoire to subsist on residual organic carbon in the natural seawater base of the medium. Thus we used unamended Pro99 medium in this set of experiments. Similar to the results with SAR11, the presence of the copiotrophic bacteria did not greatly influence the growth of *Prochlorococcus* cultures during log phase. The presence of *Marinobacter* MIT1353 did, however, increase the growth rate of *Prochlorococcus* MIT1314 slightly (from 0.64 d^−1^ to 0.68 d^−1^) and the presence of all three copiotrophs resulted in slightly higher maximum densities of MIT1314 relative to growth alone (Fig. S5). In addition, *Prochlorococcus* MIT1327 was somewhat rescued from declining cell numbers in death phase by all three of the copiotrophic strains (Fig. S5). Similar enhancements have been observed before [61,62] and are usually attributed to the ability of catalase-containing bacteria to detoxify reactive oxygen species in their surrounding environment [26,63]. This mechanism may also play a role here, as all three copiotrophic strains possess the gene necessary for catalase production. However, SAR11 HTCC7211 also possesses a catalase gene, yet it had no effect on *Prochlorococcus* growth (Fig. S4). It is unclear why, but copiotrophic catalase mutants have also been shown to rescue the growth of *Prochlorococcus* from low densities, and the addition of exogenous catalase alone cannot replicate co-culture responses [64]. Furthermore, transcriptional studies of co-cultures suggest interactions between *Prochlorococcus* and copiotrophic bacteria beyond those related to oxidative stress [65,66]. Our results support the notion that diverse copiotrophic bacteria facilitate *Prochlorococcus* growth and that these benefits are due in part to factors other than the production of catalase.

The response of the three copiotrophic bacteria to the presence of *Prochlorococcus* was strikingly different from that of SAR11, especially given that there are no added organic compounds in Pro99 medium. All three strains had an initial phase of rapid logarithmic growth (growth rate = 5.0 - 7.7 d^−1^; about an order of magnitude faster than that of SAR11), whether alone or in the presence of *Prochlorococcus*, presumably consuming organic matter present in the natural seawater base of the medium (Fig. 5). *Thalassospira* MIT1351 and *Roseobacter* MIT1352 reached higher cell abundances before entering stationary phase when grown with *Prochlorococcus* compared to growth alone (Fig. 5a-d), while this was not the case for *Marinobacter sp.* MIT1353 (Fig. 5e,f). In stark contrast to the SAR11/*Prochlorococcus* co-cultures, all three copiotrophs pulled out of stationary phase and resumed growth (at a rate of 0.49 – 0.82 d^−1^) when *Prochlorococcus* populations entered stationary phase (Fig. 5 shaded regions), displaying a diauxic growth pattern. This suggests that conditions produced by *Prochlorococcus* that are toxic to an oligotroph (e.g. SAR11) are ideal for opportunistic bacteria such as these, with more diverse functional repertoires. Comparative genomic analysis reveals that all three copiotrophic bacteria possess more genes related to sugar catabolism, hydrolysis and beta-oxidation of fatty acids, binding of extracellular solutes, and transcriptional regulation than SAR11 HTCC7211, highlighting the contrasting degrees of metabolic flexibility among these strains (Table S2).

**Figure 5.**
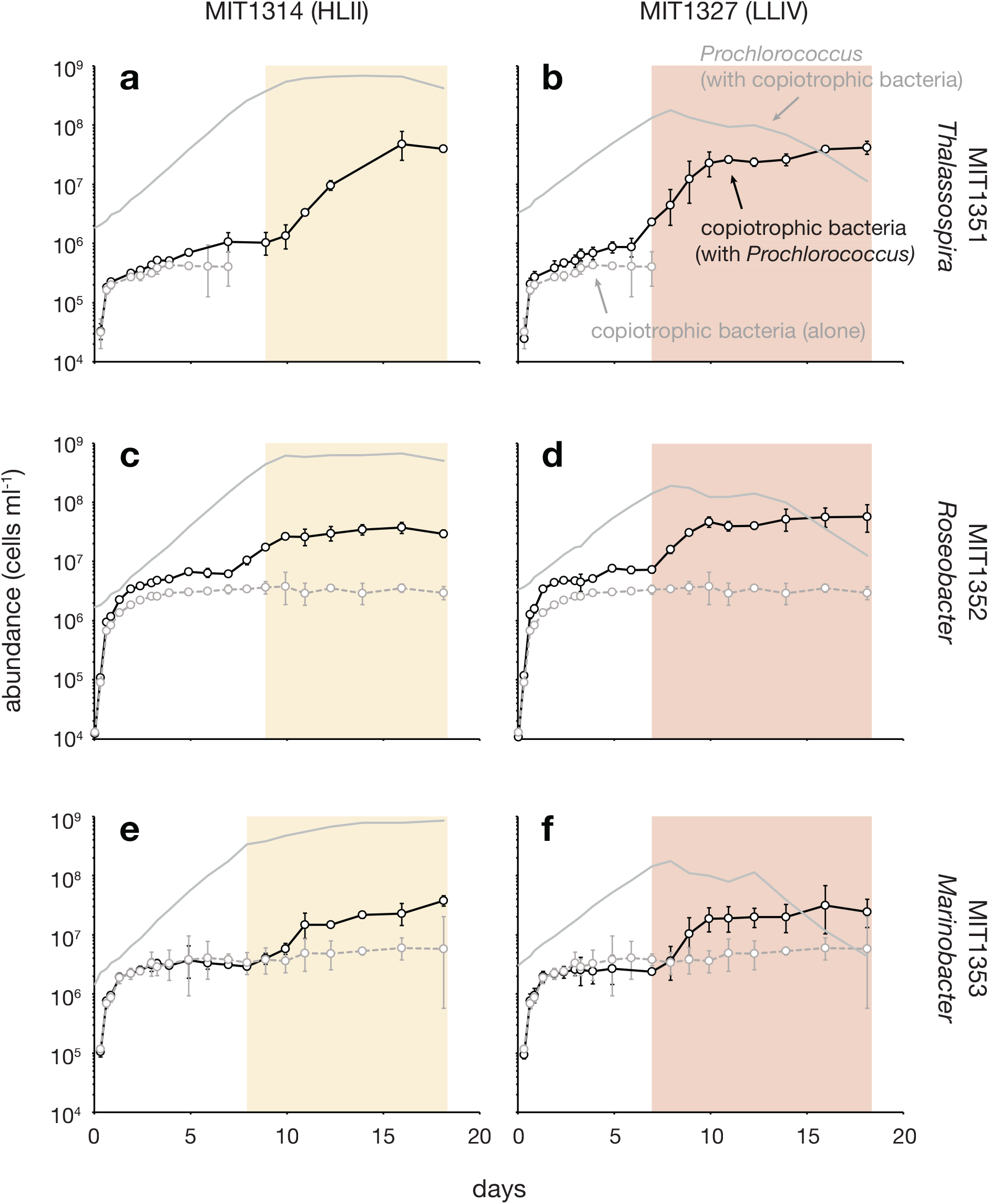
Growth of *Prochlorococcus* (solid gray lines, reproduced from Fig. S5) and copiotrophic bacteria (black lines) in co-culture compared to the growth of copiotrophic bacterial monocultures (dashed gray lines). Growth curves for *Prochlorococcus* monocultures are presented in Fig. S5. Shaded regions denote the interval when *Prochlorococcus* are in stationary phase. *Thalassospira* MIT1351 monocultures were indistinguishable from instrument noise after day 7, precluding reliable determination of cell concentrations. Circles represent the mean (± s.d.) of biological triplicates. Error bars are smaller than the size of the symbols where not visible.

As mentioned above, a caveat in comparing these two sets of experiments is that the co-culture medium used in the SAR11 experiments was, by necessity, augmented with pyruvate, glycine, methionine, and pico-to nanomolar concentrations of 9 vitamins (Table S1), whereas the medium in the copiotroph experiments was unamended. Despite this difference, we believe the comparison is illuminating, given that the organic additions cannot explain the SAR11 decline when *Prochlorococcus* enters stationary phase. Furthermore, the removal of pyruvate and glycine had no effect on this pattern (data not shown).

### *Prochlorococcus* can fulfill the central carbon requirement of SAR11

Low molecular weight organic acids, including pyruvate, lactate, oxaloacetate, acetate, and taurine are central carbon sources for SAR11, including non-glycolytic strains like HTCC7211 [19,28]. To determine if *Prochlorococcus* can provide a central carbon source to SAR11, we grew the latter with two strains of *Prochlorococcus* –MED4 (HLI clade) and MIT9313 (LLIV clade) – in a medium with no added pyruvate (ProMC). Pyruvate is thought to be an essential central carbon compound available to all SAR11 [52,67] and it was removed in order to drive the system toward pyruvate starvation. To ensure pyruvate limitation we increased the concentration of all other organic compound additions in ProMC (i.e. glycine, methionine, and 9 vitamins) 50-fold above ProMS levels, matching concentrations used to achieve maximum SAR11 densities [28]. A treatment with pyruvate added (50 μM) to ProMC served as a positive control. While the maximum abundance of SAR11 was reduced nearly 50-fold compared to the positive control when grown alone in ProMC, it was reduced only 7-fold and 9-fold respectively in co-culture with MIT9313 and MED4. This could not be attributed to pyruvate carry over, as these results were obtained after extended semi-continuous culturing in ProMC over several months to remove any residual pyruvate (Fig. 6a). We conclude that both strains of *Prochlorococcus* produce and release organic matter that fulfills the central carbon requirement of SAR11.

**Figure 6.**
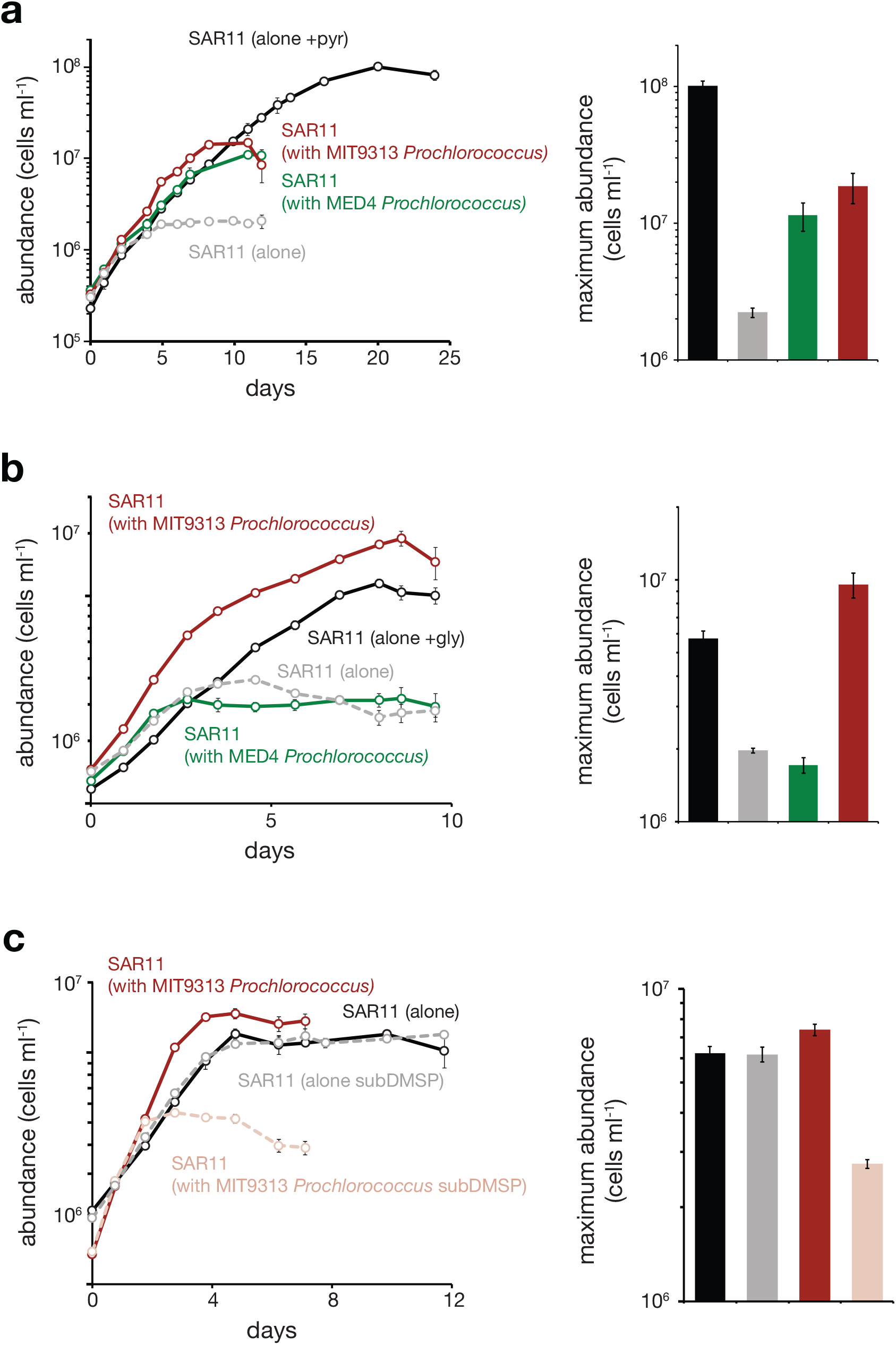
Growth of SAR11 (*Pelagibacterales sp.* HTCC7211) monocultures in the presence (black lines) and absence (dashed gray lines) of pyruvate (a) and glycine (b) compared to growth of SAR11 in co-culture with *Prochlorococcus* strains MED4 (green lines) and MIT9313 (red lines) in the absence of pyruvate (a) and glycine (b). Panel c shows the growth of SAR11 monocultures in the presence of methionine (black line) or 3-dimethylsulfoniopropionate (DMSP) (dashed gray line) compared to growth of SAR11 in co-culture with MIT9313 in the presence of methionine (red line) or DMSP (dashed pink line). Bar graphs on the right depict the maximum cell density obtained for each treatment. Bars are colored to match lines. Circles and bars represent the mean (± s.d.) of biological triplicates. Error bars are smaller than the size of the symbols where not visible.

What is *Prochlorococcus* supplying to SAR11 in these experiments? There are several candidate compounds. *Prochlorococcus* cells have been shown to release 9 to 24% of the inorganic carbon they assimilate as dissolved organic carbon, 4 to 20% of which is thought to be in the form of low molecular weight carboxylic acids, including glycolate, acetate and perhaps lactate [2]. Acetate and lactate can replace pyruvate for SAR11 growth [19]. Pyruvate is also predicted to be exported by *Prochlorococcus* as part of a strategy to maintain the cell’s redox balance [29]. Based solely on the SAR11 abundance data, it appears that *Prochlorococcus* MIT9313 provides additional labile central carbon substrates to SAR11on a per cell basis compared to MED4.

### *Prochlorococcus* MIT9313, but not MED4, can meet the glycine requirement of SAR11

SAR11 cells lack canonical genes for the biosynthesis of serine and glycine and instead rely on an exogenous source of these amino acids or their precursors [18]. To test if *Prochlorococcus* cells can meet this unique SAR11 requirement, we removed glycine from the ProMS medium (ProMS -gly) and monitored the growth of SAR11 monocultures and co-cultures with *Prochlorococcus* strains MED4 or MIT9313 after several months of semi-continuous culture to remove any traces of glycine carryover. While SAR11 monocultures were able to grow in the ProMS -gly medium - indicating that the Sargasso seawater base supplied a source of glycine, serine, or precursors of these amino acids - its maximum abundance was 4-fold lower than when grown in monoculture with 1 μM glycine. Co-culture with MED4 had no effect on this low yield, indicating that it did not provide a source of glycine or glycine substitutes to SAR11 (Fig. 6b). When in co-culture with MIT9313 however, the maximum abundance of SAR11 equaled or slightly surpassed that observed in monocultures with 1 μM glycine added. Thus, MIT9313, but not MED4, can fulfill the glycine requirement of SAR11.

It was initially surprising to us that MED4 was unable to meet SAR11’s glycine requirement, given that glycolate has been shown to substitute for glycine in SAR11 cultures [28] and glycolate production has been reported for MED4 in quantities that should have been sufficient to show an effect in our experiments [2]. The strain of SAR11 used by Carini et al (2013) was HTCC1062 (Ia.1 clade) however, and that used in our studies was HTCC7211 (Ia.3 clade). As it turns out, while HTCC1062 has the genes necessary to transport glycolate into the cell (*SAR11_0274*) and convert it to glyoxylate (*glcDEF*), a precursor of glycine [28], HTCC7211 does not, thus explaining why glycolate production by MED4 could not replace glycine in our experiments.

The most likely compound driving these disparate responses is the osmolyte glycine betaine, which has also been shown to substitute for glycine in SAR11 HTCC1062 [18,28]. SAR11 HTCC7211, the strain used in our experiments, has all the genes necessary for the uptake of glycine betaine and its conversion to glycine [68]. *Prochlorococcus* MIT9313, and other members of its LLIV clade, have the genes responsible for the biosynthesis of glycine betaine (*gbmt1/2*), along with three genes that encode an ABC transporter for this molecule (*proVWX*) [69]. Accumulation of glycine betaine has been reported in MIT9313 [70] and a recent study has shown that this accumulation can be quite significant – up to 20% of the cell’s dry weight (K. Longnecker, E. Kujawinski, personal communication). Production of glycine betaine by *Prochlorococcus* is restricted to the LLIV clade and is thus absent in MED4, further supporting the hypothesis that glycine betaine production by LLIV *Prochlorococcus* cells can fulfill the glycine requirement for SAR11 growth.

### *Prochlorococcus* and the reduced sulfur requirement of SAR11

SAR11 cells do not contain the full complement of genes necessary for assimilatory sulfate reduction, thus they rely on the production and release of reduced organic sulfur compounds such as methionine, methanethiol, or 3-dimethylsulfoniopropionate (DMSP) by other microbes for their survival [17,71]. To determine if *Prochlorococcus* could fulfill this requirement, we eliminated methionine – the only source of reduced sulfur – from the medium (ProMS -met). Neither strain of *Prochlorococcus* tested (MED4 and MIT9313) could meet SAR11’s need for reduced sulfur (data not shown).

During the initial transfers of this experiment, when the cells were growing on residual methionine, we noticed that the maximum abundance of SAR11 was 1.5-fold lower when co-cultured with *Prochlorococcus* MIT9313 (LLIV clade) vs. growth alone or with *Prochlorococcus* MED4 (HLI clade). Similar results were obtained with another *Prochlorococcus* strain from the LLIV clade (MIT1320, data not shown), suggesting that the LLIV strains may have been competing with SAR11 for a source of reduced sulfur in the Sargasso seawater medium base. Indeed, *Prochlorococcus* populations in the wild have been shown to take up DMSP [72] and, consistent with our observations, the genes required for the transport of DMSP (*proVWX*; the same transporters used for glycine betaine) [69,73,74] are found only in members of the LLIV clade of *Prochlorococcus*. To directly address the possibility of competition, we grew SAR11 alone and in co-culture with MIT9313 in a version of ProMS in which the methionine was replaced with an equimolar concentration of DMSP. SAR11 maximal abundances in co-culture with *Prochlorococcus* MIT9313 were 2-fold lower than those in monoculture, providing indirect evidence that MIT9313 may be consuming DMSP in the seawater background, and therefore compete with SAR11 for this reduced sulfur source (Fig. 6c). Thus, not only can the *Prochlorococcus* strains tested not satisfy the reduced sulfur requirement of SAR11, but it is possible that LLIV *Prochlorococcus* may be competing for some forms, such as DMSP, in the wild.

### Summary and Concluding Remarks

SAR11 grew faster in co-culture with *Prochlorococcus* than in monoculture, while the growth of *Prochlorococcus* was largely unaffected, indicating the production and release of growth factors by *Prochlorococcus* and a commensal relationship between these organisms under the conditions tested. This relationship became abruptly amensal when *Prochlorococcus* cells entered stationary phase causing rapid cell death in SAR11. That stationary phase *Prochlorococcus* cells instead triggered a secondary logarithmic growth phase in diverse copiotrophic bacteria highlights the disparate metabolic capabilities of oligotrophic and copiotrophic marine bacteria and calls for further mechanistic studies. Similarly, the variable growth response of SAR11 in co-culture with *Prochlorococcus* strains belonging to different clades demonstrates the complexity and taxonomic specificity of potential interactions in these autotroph/heterotroph pairings. Furthermore, *Prochlorococcus* MIT9313 enhanced SAR11 growth in co-culture under pyruvate and glycine limited conditions, but had the opposite effect when methionine was limiting, highlighting yet another layer in the complexity of this simple co-culture system.

Collectively, our experimental findings reveal the clade-specific ability of *Prochlorococcus* to fulfill some of the unique metabolic requirements of SAR11 and our biogeographic analyses reinforce the notion that these are two of the most ubiquitous and numerically abundant marine bacteria on Earth. Now that conditions for the steady state co-culturing of these two iconic marine microbes are established, follow up studies tracking their transcriptomes, proteomes and metabolomes will help identify the chemical exchanges and physiological dependencies that define their interactions.

## Acknowledgements

We thank Stephen Giovannoni and Amy Carter for preparation and shipment of SAR11 strain HTCC7211 and Jessica Berta-Thompson and Kristen LeGault for assisting in the isolation and purification of *Prochlorococcus* strain MIT1314. We also thank Thomas Hackl, Steven Biller, Paul Berube, Brandon Satinsky and Rogier Braakman for helpful discussions. This work was supported in part by grants from the National Science Foundation (OCE-1153588 and DBI-0424599 to S.W.C.), the Simons Foundation (Life Sciences Project Award ID 337262, S.W.C; SCOPE Award ID 329108, S.W.C), and the Gordon and Betty Moore Foundation (Grant IDs GBMF495 and GBMF4511 to S.W.C.). This paper is a contribution from the Simons Collaboration on Ocean Processes and Ecology (SCOPE) and from the NSF Center for Microbial Oceanography: Research and Education (C-MORE).

## Author contributions

J.W.B. designed and performed experiments, analyzed data and wrote the manuscript with contributions from all authors; S.L.G. performed bioinformatic analyses and writing related to biogeography; K.R. performed experiments involving copiotrophic bacteria; S.W.C. designed experiments and supervised the project.

## Supplementary Methods

### Isolation of sympatric *Prochlorococcus* and heterotrophic bacteria

We had hoped to isolate sympatric *Prochlorococcus* and oligotrophic (k-selected) heterotrophic pairs (e.g. SAR11), but we were unsuccessful. We succeeded, however, in isolating new *Prochlorococcus* strains, along with sympatric copiotrophic bacteria, from the N. Pacific (Table 1). The latter were chosen to represent a broad phylogenetic range of heterotrophic bacteria, including both alpha -and gammaproteobacteria. *Prochlorococcus* MIT1314 (HLII clade) and MIT1327 (LLIV clade) were isolated using established enrichment protocols [1] with some modifications [2], and rendered axenic via high-throughput dilution-to-extinction culturing as previously described [3]. The purity of these isolates was confirmed by flow cytometry and a suite of purity test broths: ProAC, ProMM and MPTB [3]. Sympatric heterotrophic bacteria strains MIT1351 (*Thalassospira*), MIT1352 (*Roseobacter*) and MIT1353 (*Marinobacter*) were isolated from the *Prochlorococcus* enrichment that generated the two new *Prochlorococcus* strains. The *Thalassospira* strain was isolated via high-throughput dilution-to-extinction culturing as previously described [4] using a sterile Sargasso seawater-based medium amended with ammonium (1 μM) and phosphate (0.1 μM). Dilution plates were maintained at 23 oC under constant illumination (3 μmol photons m^−2^ s^−1^). The *Roseobacter* and *Marinobacter* strains were isolated as colonies on soft agar (0.3%) pour plates consisting of Pro99 medium amended with pyruvate (0.05%) and TAPS buffer (3.75 mM). Agar plates were maintained at 23 oC under constant illumination (1 to 6 μmol photons m^−2^ s^−1^). Colonies with distinct morphologies were picked and passed at least 3 times before transfer into liquid Pro99 medium. Purity of the heterotrophic bacteria strains was assessed by flow cytometry and genomic sequencing. The strains were maintained on Sargasso seawater-based Pro99 medium in acid-washed autoclaved borosilicate glass tubes at 23 °C under constant illumination (11 μmol photons m-2 s^−1^) prior to co-culturing as, unlike SAR11, they could grow in unamended Pro99.

### Genome sequencing

Genomes of *Prochlorococcus* MIT1314 and heterotrophic bacteria strains MIT1351, MIT1352 and MIT1353 were sequenced from genomic DNA obtained from 35 to 50 ml laboratory cultures. Cells were concentrated by centrifugation (10 000 *g* for 10 min) and genomic DNA was isolated using the MasterPure complete DNA and RNA purification kit (Epicentre). 2 μg of high-molecular weight genomic DNA was sheared, and 625 – 650 ng of sheared DNA was used to construct libraries using a standard SMRTbell Template Prep Kit 1.0 and Sequencing Primer v3 (Pacific Biosciences) with the addition of a gap repair and end polishing step to ensure closed single-stranded template. Quality control analysis was performed before and after library construction using a Q-bit, NanoVue, and an Advanced Analytics DNA Fragment Analyzer, and DNA was quantified using PicoGreen assays. The average insert range for completed libraries was 21 – 28 kb. DNA libraries were sequenced using P6 chemistry on a Pacific Biosystems RS II instrument with a 240 min data collection time per SMRTcell. Data processing using SMRT Analysis 2.0 was performed to remove linkers and trim rear ends. *novo* assembly into a single contig was performed using the SMRT Portal software and the RS_HGAP_Assembly.2 protocol. The mean read quality score for each assembly was between 0.85 and 0.86 and coverage ranged from 73X to 168X. Genomes were circularized using the Geneious sequence analysis package (V7.1, Biomatters) prior to polishing using Quiver and the RS_Resequencing.1 protocol within the SMRT Portal software to obtain more accurate consensus calls. Closed circular genomes were deposited in the Joint Genome Institute’s Integrated Microbial Genomes (IMG) system and annotated using the IMG Annotation Pipeline version 4 [5,6]. Library construction and sequencing was performed at the University of Massachusetts Medical School Deep Sequencing and Molecular Biology Core Laboratories.

### Data sources

Reference genomes used in this study were downloaded as nucleotide assemblies from NCBI GenBank or Refseq and the Prokka tool [7] was used to call genes and produce annotations for all genomes. We constructed a custom nonredundant 56 million sequence reference database of microbial proteins for use with the protein homology classifier Kaiju [8]. To ensure maximum representation of marine bacteria, archaea, and microbial eukaryotes, we included translated genes/transcripts from 5 397 representative “specI” species clusters from the proGenomes database [9]; 113 transcriptomes from the Marine Microbial Eukaryote Transcriptome Sequencing Project (MMETSP) [10]; 10 509 metagenome assembled genomes from the Tara Oceans expedition [11,12], the Red Sea [13], the Baltic Sea [14], and other aquatic and terrestrial sources [15]; 994 isolate genomes from the Genomic Encyclopedia of Bacteria and Archaea [16]; 7 492 viral genomes from NCBI RefSeq [17]; 786 bacterial and archaeal genomes from MarRef [18]; and 677 marine single cell genomes [19]. Metagenomic read files to be annotated from the Tara Oceans project [20-22] were downloaded from the European Nucleotide Archive. The remaining 606 metagenomes were recently described [23] and are available from NCBI.

### Metagenome quality control

We preprocessed raw metagenome reads using the bbtools software suite (BBMap V37.90; [24]). Briefly, we trimmed reads of Illumina adapters using kmer matches (k=23) to known adapter sequences from the 5’ end of the read. We required a Maximum Hamming distance for reference kmers of 1, and we discarded reads with 3 or more ‘Ns’ or with an average quality score of less than Q20. We further trimmed overlapping reads based on insert size (tbo=t) if adapter kmers were not identified and both reads were trimmed to the minimum length of the read pair. Finally, we discarded any reads shorter than 60 bp after all adapter trimming steps. We then discarded reads containing common sequencing artifacts using kmer matches (k=31) to known Illumina sequence artifacts provided in the bbtools software suite. Reads were merged using the bbmerge tool with default parameters, a minimum insert size of 35 bp, and a minimum overlapping sequence of 12 bp. We retained all unmerged and orphaned reads that passed quality control for downstream annotation and analysis.

### Genome quality control

Because our goal was to annotate *Prochlorococcus* and SAR11 reads from shotgun metagenomes, we needed to ensure taxonomic fidelity of all genomes used as annotation references. We determined an initial genome taxonomy for all genome bins used to construct our reference genome database using checkM (V1.0.11) with the default lineage workflow [25] and ensured that all genome bins met the completion/contamination thresholds outlined in prior studies [12,15]. We then selected genomes with a taxonomic assignment of *Prochlorococcus* or SAR11 (*Pelagibacterales* order) for further analysis and confirmed taxonomic assignments using blast matches to known *Prochlorococcus* ITS sequences and by matching 16S sequences of SAR11 to the SILVA database [26]. To refine our estimates of completeness/contamination of *Prochlorococcus* genome bins, we created a custom set of 503 single copy genes from closed, isolate-derived *Prochlorococcus* genomes [27] for quality assessments with checkM. For SAR11, we used the checkM taxonomic-specific workflow with the order *Pelagibacterales*. After the custom checkM quality control, we excluded genomes from downstream phylogenetic analysis that had an estimated quality less than 30, defined as %completeness – 5 X %contamination.

### Metagenome read classification

We taxonomically classified metagenome reads using the short read classifier Kaiju v1.6.0 [8] with a custom, curated reference database comprised of approximately 26 000 bacterial, archaeal, viral, and eukaryotic proteomes (methods section “Data Sources”). The taxonomic composition of the database was intended to predominantly reflect that of the marine environment, while minimizing (but not excluding) the representation of clinical, industrial, and terrestrial host-associated samples, which greatly increased the classification speed of the more than 800 metagenomes analyzed in this study. We excluded sequences from our custom database with lengths less than 20 and greater than 20 000 amino acids, removed non-standard amino acid residues, and condensed redundant protein sequences to a single representative sequence to which we assigned a lowest common ancestor taxonomy identifier from the NCBI taxonomy database [28]. We used Kaiju in ‘mem’ mode with a minimum BLOSUM62 score of 65 for fragment matching and a SEG filter to mask low complexity sequences. We allowed for up to three amino acid substitutions in each match alignment and excluded matches with a Kaiju “expect value” less than 0.05. Using this curated database, the majority of reads in our study (54% ± 10%) could be classified across all metagenomes.

### Relative taxonomic abundance from total reads and marker genes

We used the abundance of single-copy, universal microbial marker genes [29-31] to estimate taxonomic abundances based on total counts of taxonomically recruited reads. In this approach, universal genes present in equal copy number in all bacteria and archaea are quantified and taxonomically classified, which addresses potential systematic biases introduced by changes in average genome size (AGS) and/or taxonomic composition. Thus, the abundance of different taxonomic groups can be expressed in terms of discrete ‘genomic equivalents’ rather than individual reads, which is independent of AGS and more intuitively related to the abundance of individual taxa.

*Prochlorococcus* (1.6-2.5 Mb) and SAR11 (1.2-1.4 Mb) both have small genomes, thus their relative abundance estimated from total recruited reads could underestimate their true abundance, while overestimating the abundance of microbes with larger genomes. Indeed, we found that relative read abundance and estimated relative genome equivalents (using MicrobeCensus V1.1.1; [32], see below) of *Prochlorococcus* and SAR11 had a strong linear relationship, while read counts alone systematically underestimated the relative abundance of either organism, especially when they constituted a quarter or more of the identifiable bacteria and archaea in a given sample (Figure S6). This finding supports previous work demonstrating that AGS differences between microbial groups can distort relative abundance estimates based on total read classification depending on the degree of variability and the relative contribution of differently sized genomes to the total genome pool in a microbial community [32,33].

### Relative abundance estimates of *Prochlorococcus* and SAR11 using genome equivalents

To address potential artifacts related to genome size, we used a hybrid marker gene/total read classification approach to estimate genome equivalents for taxonomically resolved read groups in each metagenome. We first classified all reads in each metagenome using Kaiju and the custom database described above, and then pulled reads mapping to *Prochlorococcus* or SAR11, and all reads annotated as bacteria and archaea, while excluding reads annotated as microbial eukaryotes, viruses, or those with no classification. We then quantified marker genes within each taxonomically partitioned read pool using the MicrobeCensus (V1.1.1) algorithm [32]. MicrobeCensus quantifies 30 well-characterized single-copy microbial marker genes using sensitive gene family-specific sequence similarity thresholds, but without any taxonomic distinction. Then the algorithm uses proportionality constants and weights for each gene family to relate counts to the relative abundance for each marker gene. This relative abundance is multiplied by the inverse of linear model coefficients for each gene family to obtain an estimate of AGS for each marker family, the weighted average of which becomes the total AGS. In each step, all model coefficients, thresholds, weights and proportionality constants have been estimated from simulated metagenomes of defined taxonomic abundance and composition.

The number of genome equivalents within a taxonomically resolved read pool is calculated by the total number of bases divided by the estimated AGS. In this way, we obtained information about AGS and the number of genome equivalents for the genus-integrated *Prochlorococcus* and order-integrated SAR11 (*Pelagibacterales*) populations in each metagenome. The mean AGS was 1.46 ± 0.24 Mb for *Prochlorococcus* and 1.2 ± 0.10 Mb for SAR11 across all samples, which agrees well with the genome sizes of sequenced isolates from these groups [27,34]. We then compared the proportion of *Prochlorococcus* or SAR11 genome equivalents to the total number of bacterial and archaeal genome equivalents detected in each sample to determine relative abundances.

The relative abundance of any taxonomic group in a metagenome can be distorted by the proportion of unclassified or unknown reads, particularly if this unclassified proportion varies randomly across samples/environments/conditions being compared. Although we constructed a taxonomically diverse and comprehensive reference database specifically tailored to the marine environment, we undoubtedly missed some of the microbial diversity present in each sample. This is a source of error endemic to any microbiome study, and metagenomic studies in under-sampled environments (e.g. marine, soil) typically have significantly lower read recruitment rates than studies in well-characterized systems (e.g. human or other host-associated). The majority of reads in our study (54% ± 10%) could be classified across all metagenomes, with the largest numbers of unclassified reads found in GEOTRACES samples, particularly in neritic zones and from latitudes greater than 30 degrees N/S. The GA10 transect in particular displayed the lowest total recruitment, as well as the lowest specific recruitment to *Prochlorococcus* and SAR11. In contrast, samples from the Tara oceans project, particularly those from equatorial waters, had the highest classification rates - often approaching 80%. This classification discrepancy between the two sequencing projects is likely because the Tara samples used here were prefiltered to exclude larger eukaryotes, which are often poorly represented in reference database [22], whereas the GEOTRACEs samples were not prefiltered [23]. Additionally, our reference database included over 2 500 metagenome assembled genomes derived from the Tara oceans prokaryotic size fractions [11,12], which we would expect to strongly recruit metagenomic reads from Tara samples.

We estimated the abundance of marker genes (using MicrobeCensus) in the unclassified read fraction (from Kaiju) for 100 randomly selected metagenomes, and found that the number of genome equivalents ‘missed’ in this unclassified fraction averaged < 5% of the total number of genome equivalents estimated for all bacteria and archaea in each metagenome, suggesting that our unclassified read pools are predominantly comprised of novel, unknown gene families rather than universal single-copy genes. Therefore, we expect our genome equivalent estimates to be robust to variability in read recruitment rates, however it is possible that our methodological approach may have missed some small fraction of highly novel bacterial/archaeal genome equivalents, resulting in a slight overestimation of the relative contribution of SAR11 and *Prochlorococcus* to the total prokaryotic community.

### Phylogenetic inference

The *Prochlorococcus* phylogeny (Figure S1a) includes 135 isolate, single-cell, and metagenome-assembled genomes (estimated > 90% complete), and the SAR11 phylogeny (Figure S1b) includes 92 isolate, single-cell, and metagenome-assembled genomes (estimated > 70% complete). These completeness thresholds were selected as a compromise between maximum completeness and phylogenetic diversity of the available genomes. Each phylogeny is based on a concatenated protein multiple sequence alignment of 120 taxonomically conserved single copy marker genes [15,35]. We used GTDB-Tk (V0.0.7;https://github.com/Ecogenomics/GTDBTk) with default settings to generate the alignments using HMMER V3.1b2; http://hmmer.org/). We pruned/trimmed both alignments to remove genomes that filled fewer than 50% of alignment positions in order to remove columns represented by fewer than 50% of all taxa and to remove columns with no single amino acid residue occurring at a frequency greater than 25%. We further trimmed the alignments using trimAl with the automated *-gappyout* option to trim columns based on their gap distribution. Genomes were pruned from a starting tree to allow a maximum average distance to closest leaf (ADCL) of 0.003 to reduce the phylogenetic redundancy within the overall phylogeny [36]. We constructed phylogenies using Maximum Likelihood inference with multithreaded RAxML (V8.2.9; [37]) using the GAMMA model of rate heterogeneity, empirically determined base frequencies, and the LG substitution model [38] (PROTGAMMALGF). Branch support is based on 250 resampled bootstrap trees.

## Supplementary Figure Legends

**Figure S1.**
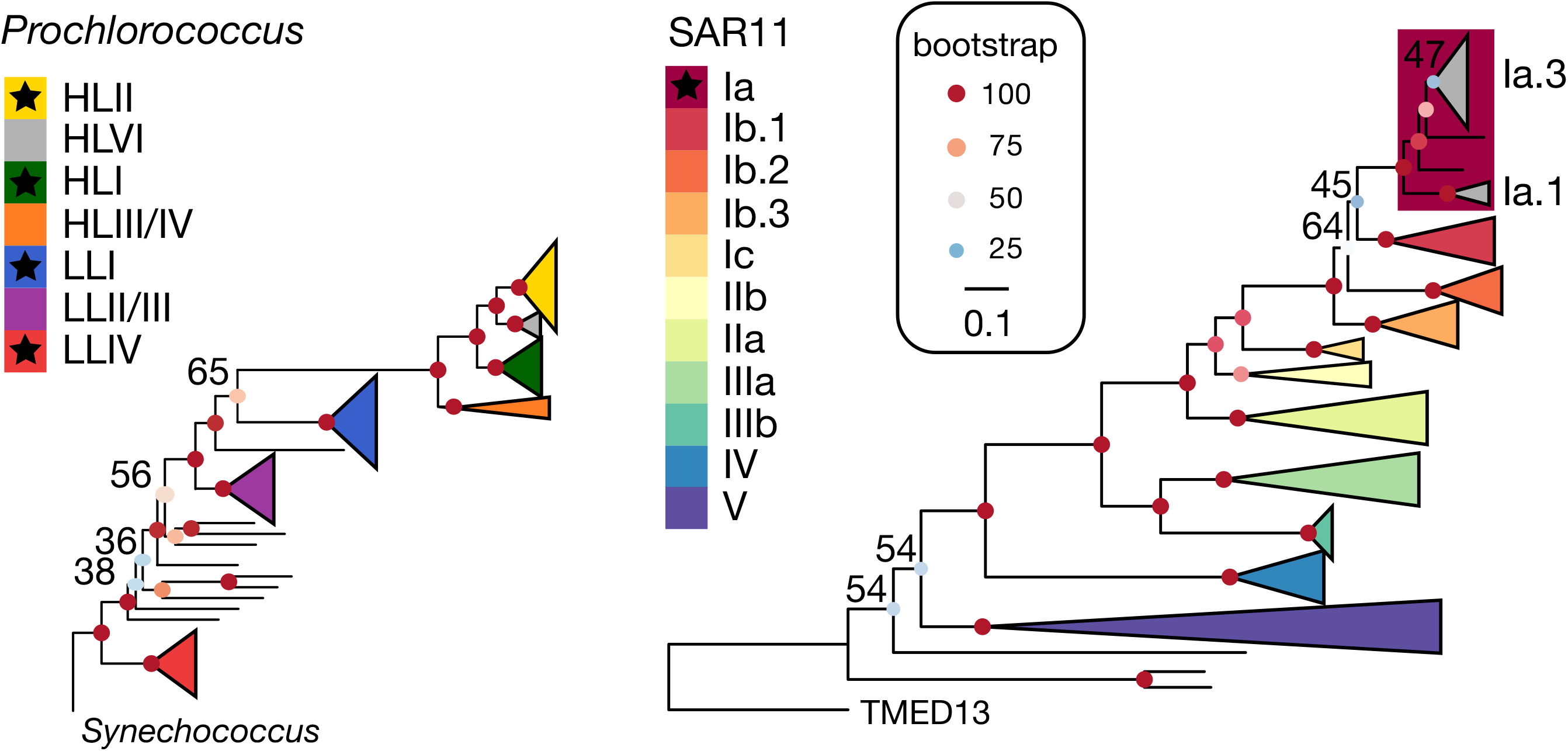
Genome phylogenies of *Prochlorococcus* and SAR11. The phylogenies include 135 *Prochlorococcus* and 92 SAR11 isolate, single-cell, and metagenome-assembled genomes. Bootstrap values are represented by filled circles and are sized and colored according to support magnitude. The scale bar is shared for both trees and represents 0.1 amino acid substitutions. The *Prochlorococcus* tree is rooted on *Synechococcus* sp. WH5701 and the SAR11 tree is rooted on TMED13, a metagenome-assembled genome from Tara Oceans distantly related to ‘*Candidatus* Pelagibacter ubique’. Monophyletic clades for both lineages are represented by colored cartoon triangles. Bootstrap support for all clades is greater than 90%, unless specifically denoted with text. Clades utilized in co-culture experiments are marked with a star. Note that the inclusion of new genomes in this study resulted in a polyphyletic origin for the canonical SAR11 clade Ib, therefore we have partitioned clade Ib into 3 monophyletic subclades (Ib.1, Ib.2 and Ib.3).

**Figure S2.**
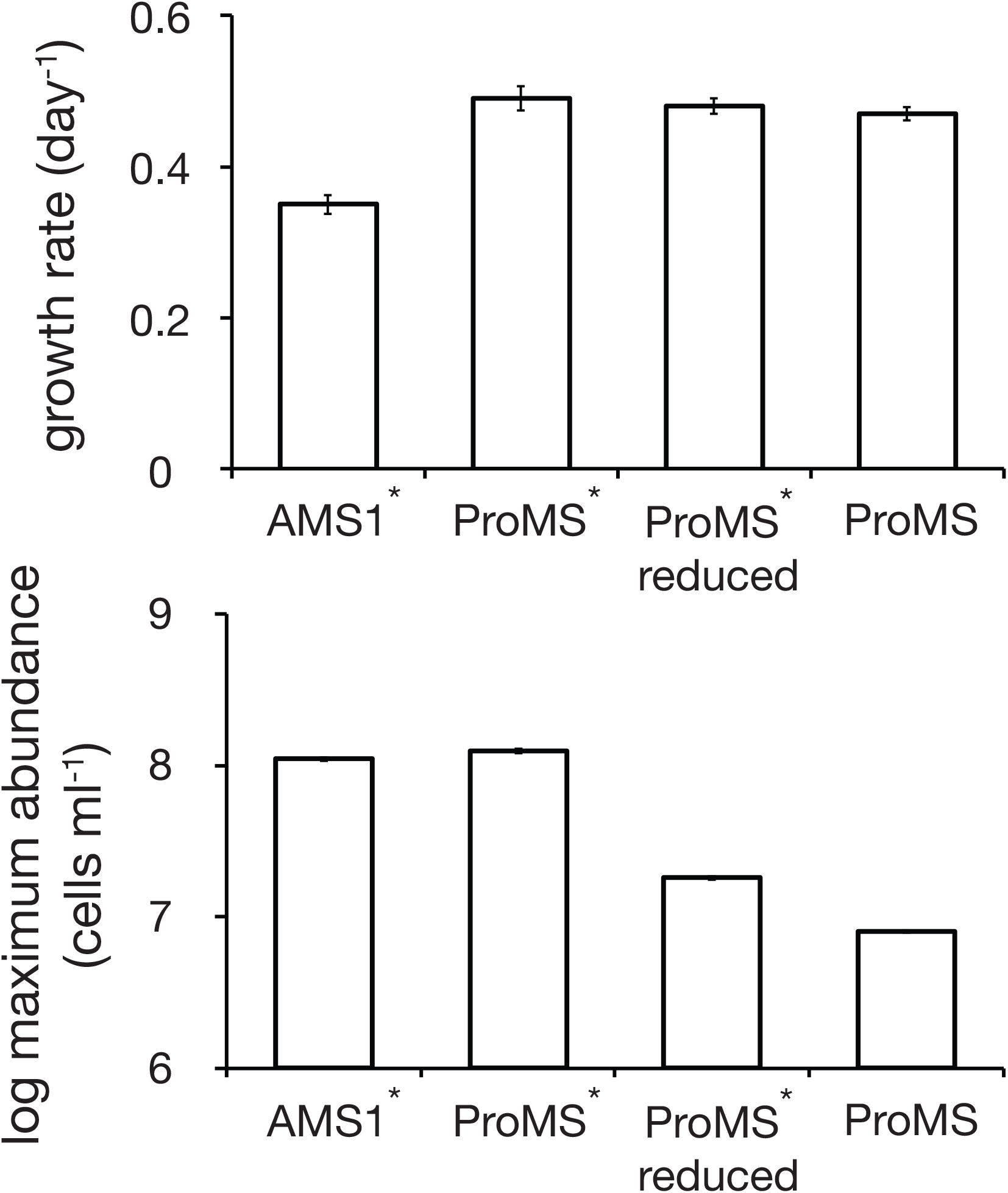
Growth rate (top) and maximum cell abundance (bottom) of SAR11 (*Pelagibacterales sp.* HTCC7211) monocultures grown in various artificial and natural seawater-based media types. AMS1* is the artificial medium designed by Carini et al. for SAR11 [39] supplemented with pyruvate (50 μM), glycine (50 μM) and methionine (10 μM). ProMS* is natural seawater based Pro99 medium [1] supplemented with pyruvate (50 μM), glycine (50 μM), methionine (10 μM) and the vitamin mix used for AMS1. ProMS* reduced is Pro99 medium supplemented with pyruvate (2.5 μM), glycine (2.5 μM), methionine (0.5 μM) and the vitamin mix used for AMS1. ProMS is the medium designed for the co-culture of *Prochlorococcus* and SAR11 (see Table S1). Bars represent the mean (± s.d.) of at least 6 consecutive semi-continuous batch cultures.

**Figure S3.**
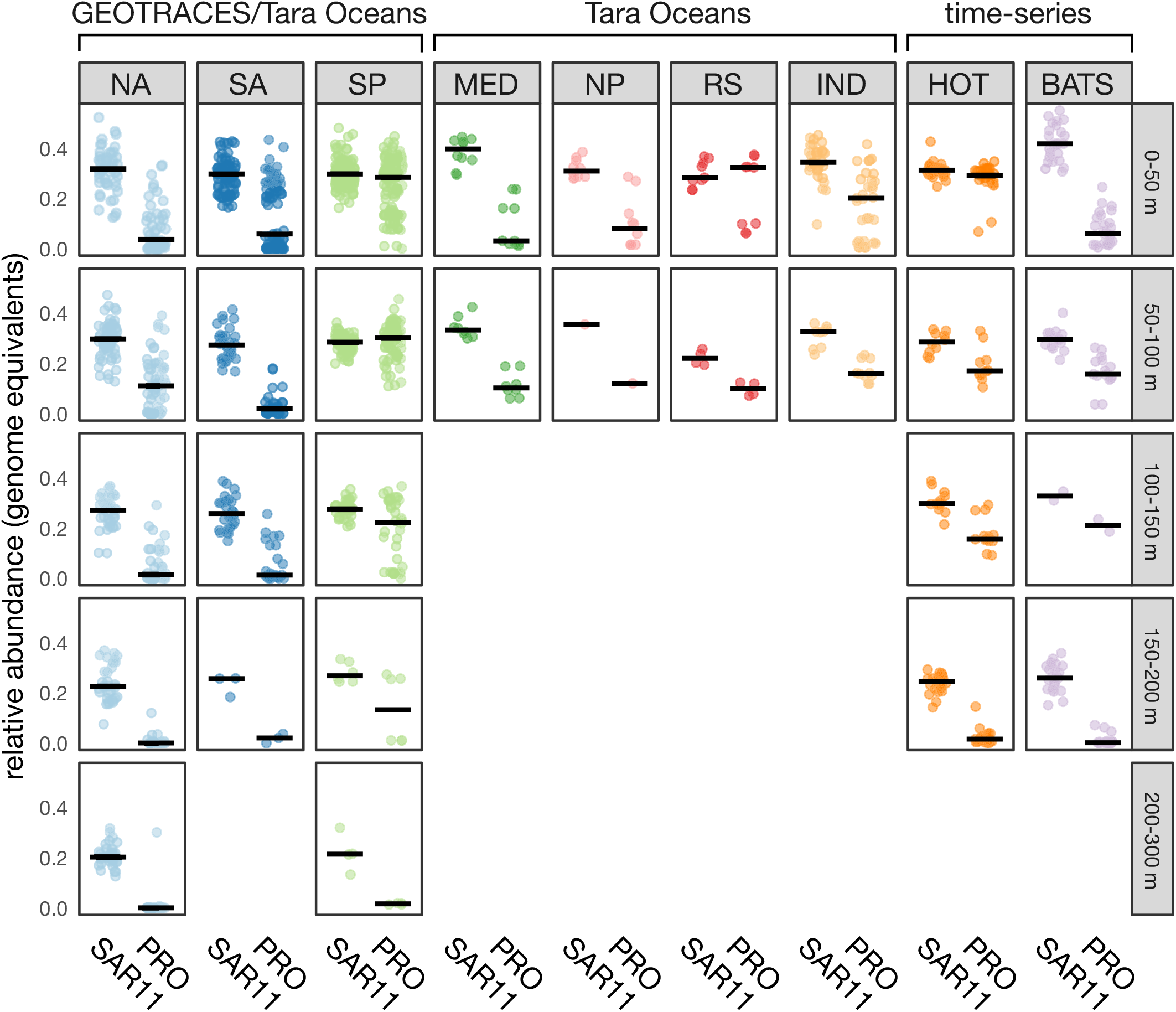
Relative abundances of *Prochlorococcus* and SAR11 estimated genome equivalents at different depths and within different ocean basins. Vertical axes represent relative abundance of genome equivalents and horizontal axes represents *Prochlorococcus* (PRO) or SAR11. Horizontal facets bins samples based on ocean region (NA=North Atlantic, SA=South Atlantic, SP=South Pacific, MED=Mediterranean, NP=North Pacific, RS=Red Sea, IND=Indian, HOT=Hawaii Ocean Time-series, BATS=Bermuda Atlantic Time-series) and vertical facets bin by depth ranges. Black horizontal bars represent the median abundance of PRO or SAR11 genome equivalents and points represent individual samples within each region/depth bin. Sample origin (GEOTRACES, Tara Oceans, or HOT and BATS Time-series) are denoted above the horizontal facets.

**Figure S4.**
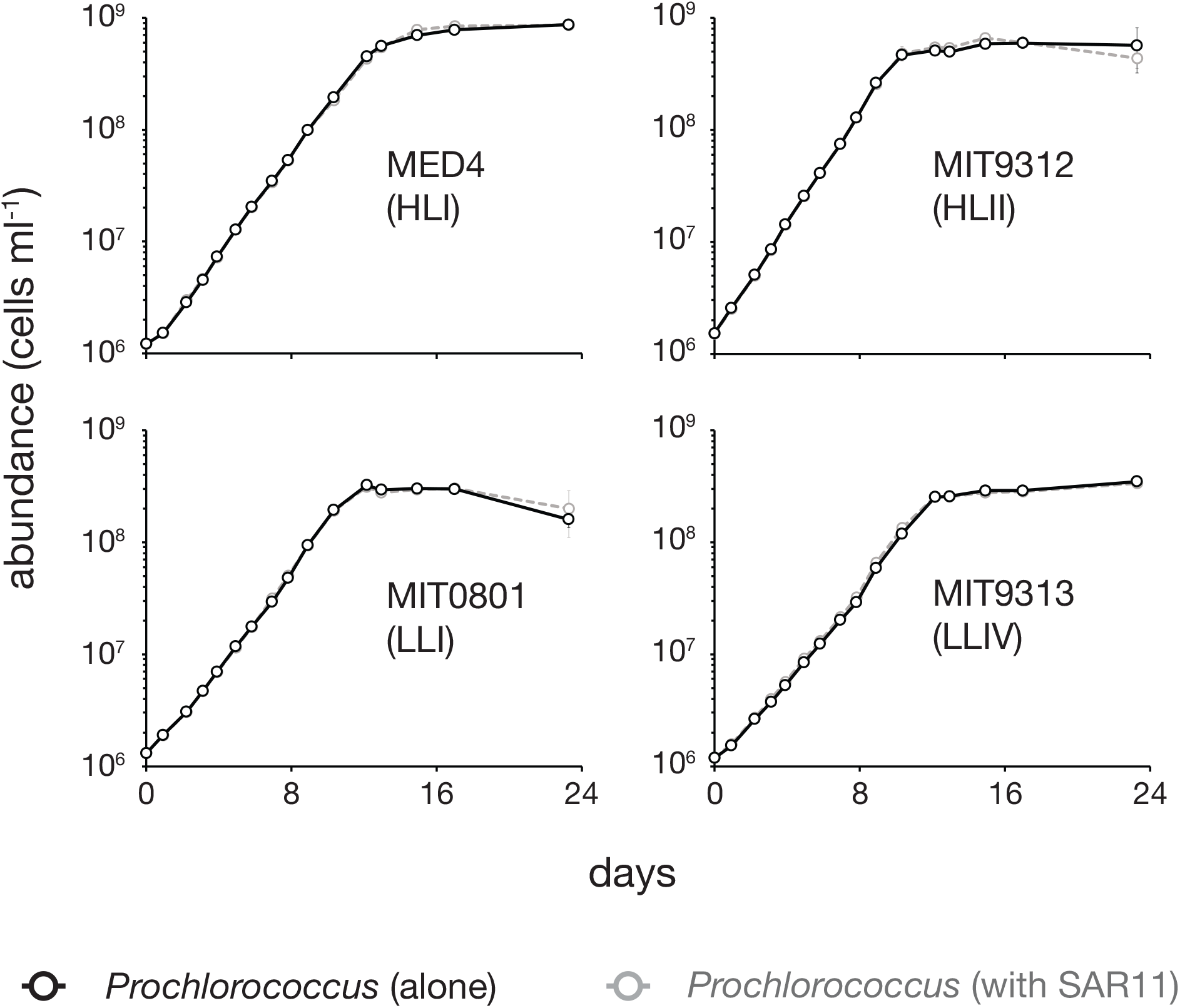
Growth of *Prochlorococcus* strains from multiple high-light (HL) and low-light (LL) adapted clades in co-culture with SAR11 (*Pelagibacterales sp.* HTCC7211) (dashed gray lines) compared to the growth of *Prochlorococcus* strains in monoculture (black lines). Open circles represent the mean (± s.d.) of biological triplicates. Error bars are smaller than the size of the symbols where not visible.

**Figure S5.**
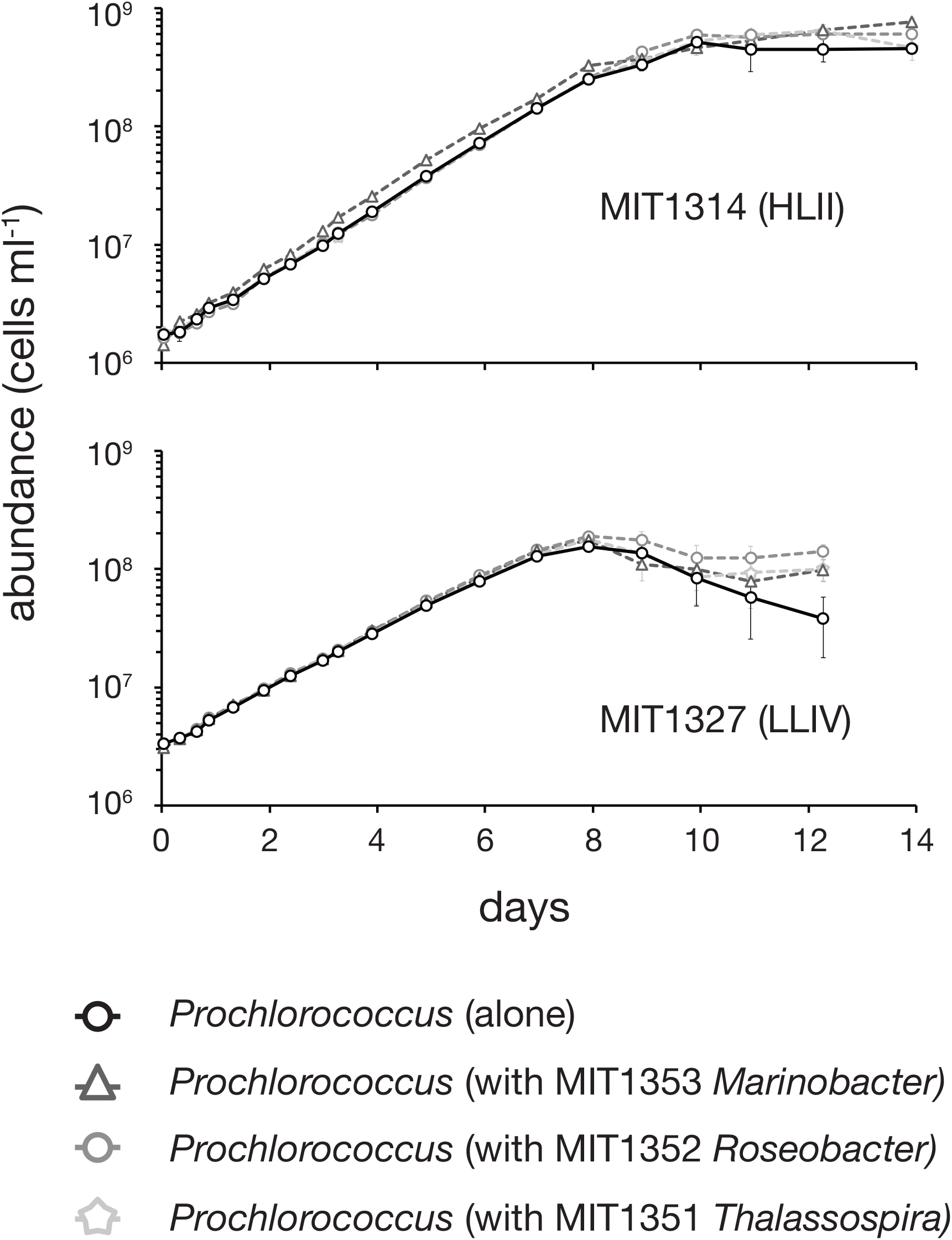
Growth of *Prochlorococcus* strains MIT1314 (HLII clade; top) and MIT1327 (LLIV clade; bottom) in co-culture with sympatric copiotrophic bacteria: *Thalassospira sp.* MIT1351 (dashed gray stars), *Roseobacter sp.* MIT1352 (dashed gray circles) and *Marinobacter sp.* MIT1353 (dashed gray triangles) compared to the growth of *Prochlorococcus* strains in monoculture (black lines). Open symbols represent the mean (± s.d.) of biological triplicates. Error bars are smaller than the size of the symbols where not visible.

**Figure S6.**
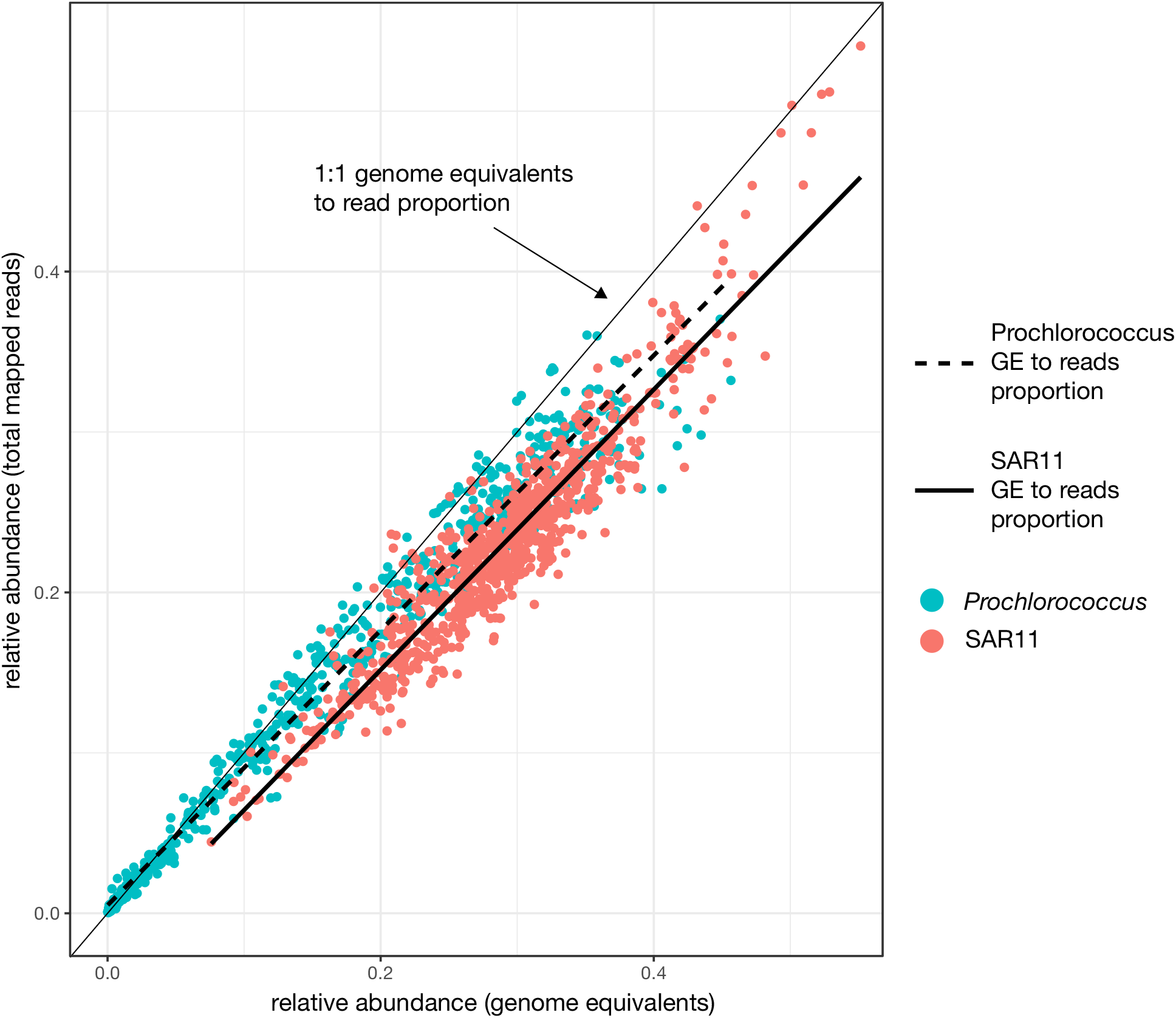
Relative abundances of *Prochlorococcus* and SAR11 based on estimated genome equivalents (GE) or based on the total fraction of mapped reads. Each point represents a metagenome sample. The thin solid black line represents the 1:1 ratio of equal proportion of mapped reads to GE. In most cases, both *Prochlorococcus* and SAR11 estimates fall below this line, indicating that relative abundances determined by total mapped reads underestimate the relative abundances of discrete genomes in each sample. Thick dashed/solid lines are linear models fit to the relative abundance of *Prochlorococcus* (dashed) and SAR11 (solid).

**Table S1.** Composition of ProMS medium for co-culture of *Prochlorococcus* and SAR11. Filter-sterilized surface seawater from the Sargasso Sea was used as the base. The medium was modified, through trial and error, from Pro99 [1] and AMS1 [39].

| Component | Final Concentration |
| --- | --- |
| *Macronutrients* |  |
| NH<sub>4</sub>Cl | 800 µM |
| NaH<sub>2</sub>PO<sub>4</sub>·H<sub>2</sub>O | 50 µM |
| *Trace Metals* |  |
| Na<sub>2</sub>EDTA·2H<sub>2</sub>O | 1.17 µM |
| FeCl<sub>3</sub>·6H<sub>2</sub>O | 1.17 µM |
| MnCl<sub>2</sub>·4H<sub>2</sub>O | 90 nM |
| ZnSO<sub>4</sub>·7H<sub>2</sub>O | 8 nM |
| CoCl<sub>2</sub>·6H<sub>2</sub>O | 5 nM |
| Na<sub>2</sub>MoO<sub>4</sub>·2H<sub>2</sub>O | 3 nM |
| Na<sub>2</sub>SeO<sub>3</sub> | 10 nM |
| NiCl<sub>2</sub>·6H<sub>2</sub>O | 10 nM |
| *Vitamins* |  |
| B<sub>1</sub> | 120 nM |
| B<sub>3</sub> | 16 nM |
| B<sub>5</sub> | 8.5 nM |
| B<sub>6</sub> | 10 nM |
| B<sub>7</sub> | 80 pM |
| B<sub>9</sub> | 80 pM |
| B<sub>12</sub> | 14 pM |
| Myo-inositol | 120 nM |
| 4-Aminobenzoic acid | 1.2 nM |
| Sodium pyruvate | 1 µM |
| Glycine | 1 µM |
| L-Methionine | 200 nM |

**Table S2.** The number of genes from selected protein families present in the genomes of the 4 heterotrophic bacteria discussed in this study. Protein families underrepresented in SAR11 (*Pelagibacterales sp.* HTCC7211) compared to the other 3 heterotrophic bacteria strains were selected for inclusion in the table.

| **Pfam ID** | **Description** | **SAR11**<br>**HTCC7211** | **Thalassospira**<br>**MIT1351** | **Roseobacter**<br>**MIT1352** | **Marinobacter**<br>**MIT1353** |
| --- | --- | --- | --- | --- | --- |
| pfam00027 | Cyclic nucleotide-binding domain | 0 | 13 | 5 | 6 |
| pfam00196 | Bacterial regulatory proteins, luxR family | 0 | 6 | 17 | 18 |
| pfam00440 | Bacterial regulatory proteins, tetR family | 0 | 24 | 21 | 22 |
| pfam00563 | EAL domain | 0 | 18 | 4 | 23 |
| pfam00970 | Oxidoreductase FAD-binding domain | 0 | 6 | 4 | 6 |
| pfam00989 | PAS fold | 0 | 8 | 6 | 11 |
| pfam00990 | Diguanylate cyclase, GGDEF domain | 0 | 32 | 7 | 50 |
| pfam01627 | Hpt domain | 0 | 5 | 4 | 6 |
| pfam07729 | FCD domain | 0 | 21 | 11 | 7 |
| pfam08448 | PAS fold | 0 | 6 | 3 | 5 |
| pfam12418 | Acyl-CoA dehydrogenase N terminal | 0 | 2 | 2 | 6 |
| pfam12625 | Arabinose-binding domain of AraC transcription regulator, N-term | 0 | 1 | 6 | 12 |
| pfam12697 | Alpha/beta hydrolase family | 0 | 4 | 9 | 6 |
| pfam12833 | Helix-turn-helix domain | 0 | 34 | 18 | 23 |
| pfam13426 | PAS domain | 0 | 16 | 9 | 9 |
| pfam00126 | Bacterial regulatory helix-turn-helix protein, lysR family | 1 | 55 | 37 | 21 |
| pfam00202 | Aminotransferase class-III | 2 | 12 | 8 | 6 |
| pfam00270 | DEAD/DEAH box helicase | 2 | 8 | 9 | 11 |
| pfam03061 | Thioesterase superfamily | 2 | 9 | 10 | 11 |
| pfam08240 | Alcohol dehydrogenase GroES-like domain | 2 | 10 | 9 | 9 |
| pfam00107 | Zinc-binding dehydrogenase | 3 | 9 | 10 | 8 |
| pfam00293 | NUDIX domain | 3 | 8 | 11 | 8 |
| pfam00378 | Enoyl-CoA hydratase/isomerase | 3 | 9 | 13 | 12 |
| pfam00571 | CBS domain | 3 | 10 | 10 | 13 |
| pfam00672 | HAMP domain | 3 | 35 | 22 | 23 |
| pfam04055 | Radical SAM superfamily | 3 | 10 | 11 | 14 |
| pfam00106 | short chain dehydrogenase | 4 | 13 | 13 | 16 |
| pfam00155 | Aminotransferase class I and II | 4 | 14 | 17 | 13 |
| pfam00512 | His Kinase A (phospho-acceptor) domain | 4 | 25 | 41 | 30 |
| pfam00497 | Bacterial extracellular solute-binding proteins, family 3 | 5 | 24 | 15 | 14 |

